# VILOCA: Sequencing quality-aware haplotype reconstruction and mutation calling for short- and long-read data

**DOI:** 10.1101/2024.06.06.597712

**Authors:** Lara Fuhrmann, Benjamin Langer, Ivan Topolsky, Niko Beerenwinkel

**Affiliations:** Department of Biosystems Science and Engineering, ETH Zurich, Basel, 4058, Switzerland; SIB Swiss Institute of Bioinformatics, Basel, 4058, Switzerland

## Abstract

RNA viruses exist in large heterogeneous populations within their host. The structure and diversity of virus populations affects disease progression and treatment outcomes. Next-generation sequencing allows detailed viral population analysis, but inferring diversity from error-prone reads is challenging. Here, we present VILOCA, a method for mutation calling and reconstruction of local haplotypes from short- and long-read viral sequencing data. Local haplotypes refer to genomic regions that have approximately the length of the input reads. VILOCA recovers local haplotypes by using a Dirichlet process mixture model to cluster reads around their unobserved haplotypes and leveraging quality scores of the sequencing reads. We assessed the performance of VILOCA in terms of mutation calling and haplotype reconstruction accuracy on simulated and experimental Illumina, PacBio, and Oxford Nanopore data. On simulated and experimental Illumina data, VILOCA performed better or similar to existing methods. On the simulated long-read data, VILOCA is able to recover on average 82% of the ground truth mutations with perfect precision compared to only 64% recall and 90% precision of the second-best method. In summary, VILOCA provides significantly improved accuracy in mutation and haplotype calling, especially for long-read sequencing data, and therefore facilitates the comprehensive characterization of heterogeneous within-host viral populations.

## 1 Introduction

RNA viruses and retroviruses, such as Human Immunodeficiency Virus (HIV), SARS-CoV-2, and Influenza A virus (IAV), exhibit distinct characteristics including high mutation rates, large population sizes, and short generation times. These traits contribute to the continuous generation of new viral haplotypes and the accumulation of genetic diversity within infected hosts [1]. Understanding this within-host viral diversity is crucial as it plays a significant role in determining virulence and treatment outcomes [2, 3]. Moreover, the identification of low-frequency mutations and haplotypes is of clinical importance as they have been associated with drug resistance and treatment failure [4].

Complementing traditional clinical sample analysis, wastewater samples have become increasingly important in pathogen surveillance. In response to the SARS-CoV-2 pandemic, numerous countries have implemented wastewater surveillance programs to monitor the emergence, spread, and distribution of different variants of concern [5, 6, 7]. Wastewater surveillance has also demonstrated its effectiveness in monitoring the circulation of poliovirus strains, playing a crucial role in optimizing eradication campaigns [8]. Additional research and development is underway to establish wastewater monitoring for other viruses, such as influenza, rota, or norovirus [9, 10, 11].

Next-generation sequencing (NGS) is a powerful tool for uncovering the complex structure of viral populations in both clinical and environmental samples. Advancements in NGS have improved its cost-efficiency and the accessibility of high-coverage viral NGS data, which allow the detection of low-frequency mutations and haplotypes. In-depth analysis of viral within-host diversity has primarily relied on Illumina reads, given their relatively low error rates (*<* 1% [12]). However, one limitation of Illumina technology is the short read length, typically up to 2 x 250 base pairs. Recent technological developments, implemented in PacBio and Oxford Nanopore sequencing, have reduced error rates in long reads. As a result, these technologies have become increasingly attractive for analyzing within-sample viral diversity, as they can generate reads with lengths exceeding 10 kilobases, enabling complete coverage of some viral genomes.

In order to distinguish true biological variation from technical errors, a variety of tools have been developed to reconstruct viral diversity from NGS data [13, 14], focusing on different spatial levels of the genome, including single-nucleotide variants (SNVs), local haplotypes, and global haplotypes (i.e., full-length genomes). While in the past, most tools focused on utilizing short reads, more recently, tools have emerged that can process long-read data [15, 16]. To enhance accuracy, some of these tools utilize the information from linked or co-occurring mutations, including CliqueSNV [16], ShoRAH [17], and PredictHaplo [18]. In a recent benchmarking study for global haplotype reconstruction [19, 20], PredictHaplo and CliqueSNV have shown good performance in settings of relatively low mutation rate [19]. However, in cases where the mutation rate is higher and consequently the viral diversity greater, such as with HIV, all methods included in the study performed rather poorly.

Here, we present VILOCA (VIral LOcal haplotype reconstruction and mutation CAlling for short and long read data), a statistical model and computational tool for single-nucleotide variant calling and local haplotype reconstruction from both short-read and long-read data. By incorporating position-wise sequencing quality scores and information on co-occurring mutations, VILOCA enhances the accuracy and precision in predicting diversity within mixed samples. We employ a finite Dirichlet Process mixture model that clusters reads according to their unobserved haplotypes of origin. Reads are assigned to the most suitable haplotype using a sequencing error process that takes into account the sequencing quality scores specific to each read. In contrast to other approaches, such as ShoRAH and PredictHaplo, which rely on sampling methods to learn the model parameters, we have developed a variational inference algorithm that utilizes a mean field approximation of the posterior distribution and assumes independence among the latent variables of the model. This implementation enhances the feasibility of processing very diverse samples by avoiding convergence problems commonly encountered with sampling methods. We comprehensively evaluate VILOCA using simulated Illumina, PacBio, and ONT data, varying the complexity of the underlying haplotype populations. In addition, we analyze three real experimental samples where the underlying ground truth viral strains are known. Finally, to illustrate VILOCA’s usefulness, we apply it to Polio and SARS-CoV-2 wastewater data, as well as to longitudinal clinical samples from HIV-positive patients. VILOCA is available at https://github.com/cbg-ethz/VILOCA.

## 2 Methods

We developed VILOCA, a computational method to reconstruct haplotypes and identify mutations based on aligned sequencing read data. It takes an alignment file and a reference sequence as input and generates local haplotypes and calls mutations, including insertions and deletions, with respect to the reference sequence. VILOCA proceeds in three main steps: (1) tiling of the genome into local regions, (2) haplotype reconstruction for each local region, and (3) postprocessing by applying a strand bias filter (Figure 1A).

**Figure 1:**
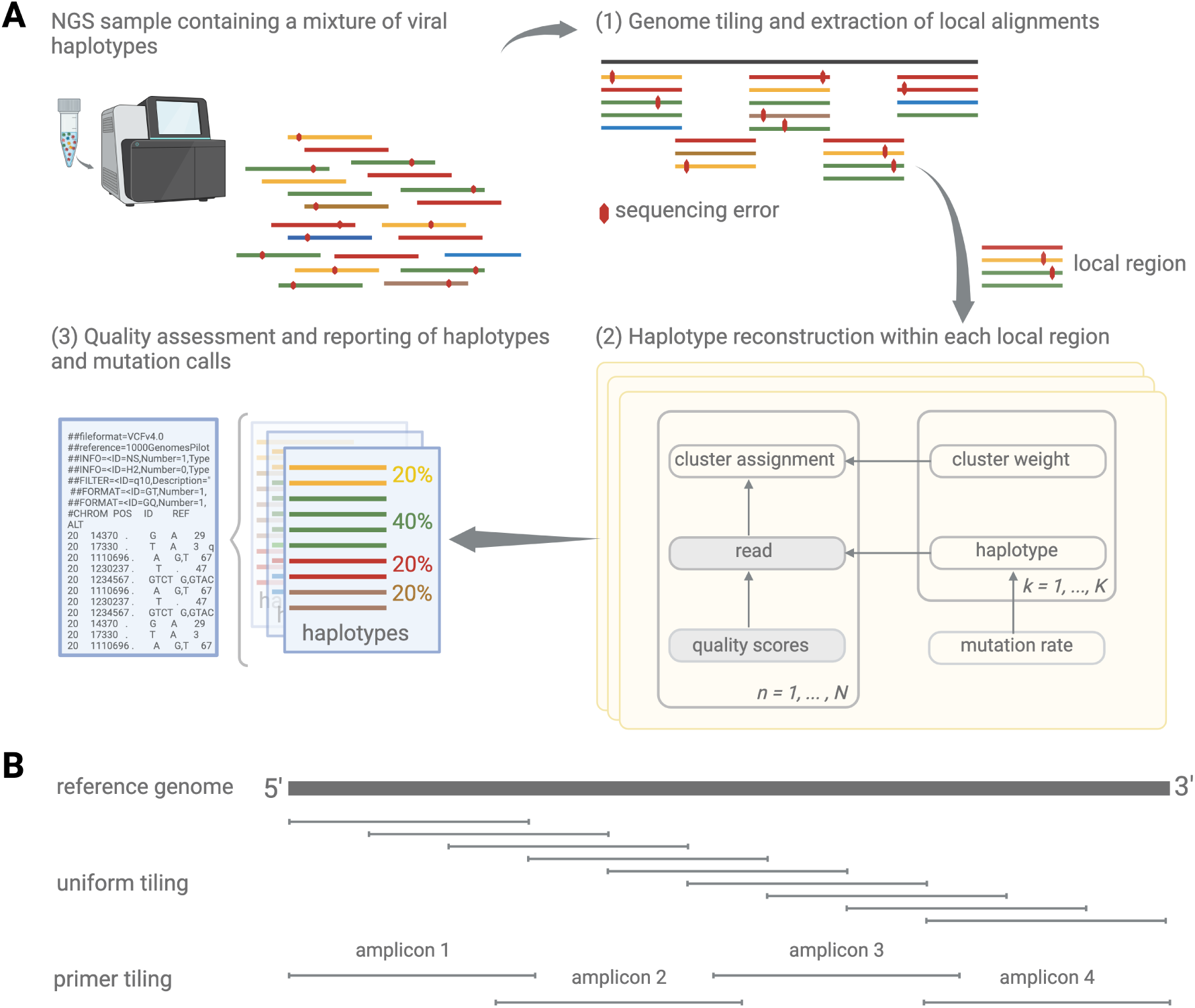
Overview of VILOCA. (A) VILOCA takes the aligned sequencing reads as input and proceeds through three main steps. (1) It defines the genome tiling and extracts the corresponding local alignments from the input alignment file. (2) For each local region, haplotypes are reconstructed using a clustering approach. (3) Finally, mutation calls across all regions are consolidated and subjected to a quality assessment. This assessment includes applying a strand bias filter for pairedend reads and evaluation of the posterior score. Subsequently, the tool reports local haplotypes and mutation calls that successfully pass the quality assessment. (B) Schematic illustration of two genome tiling approaches. In the case of uniform genome tiling, a sliding window method is utilized to distribute local regions of uniform length across the genome. Each position is covered by a number of user-defined local regions. Alternatively, when the primer strategy is known or can be specified, a bed file outlining the amplicons by start and end positions is used to define the regions.

### Genome tiling

In the first step, VILOCA divides the genome into local regions. This tiling depends on the read length and the read distribution along the genome. There are two options for users to define the genome tiling: the uniform tiling mode and the primer tiling mode (Figure 1B). The uniform tiling mode is recommended if reads are uniformly distributed and coverage is approximately even along the genome. In this mode, VILOCA divides the genome into overlapping local regions of equal length which is approximately equal to the input read length (uniform tiling, Figure 1B). Each position is covered by a user-provided number of local regions. Alternatively, users can provide a bed-file defining the local regions to run VILOCA in primer tiling mode. This is especially useful if the primer protocol used for library preparation is known, as the amplicon positions can be used to define the genome tiling (primer tiling, Figure 1B). In the case of long-read sequencing data, where reads cover the full genome, a single local region can be selected, namely the full genome, effectively enabling global haplotype reconstruction in this setting.

After defining the tiling, local alignments are created for each region by fetching reads that cover the specific region. A read is included in the local alignment if it covers the region by at least the user-defined ”win_min_ext” threshold (default: 0.85). Any segments within the region that are not covered by the read are filled with Ns. In general, the genome tiling is always a trade-off between maximizing the per-position coverage and the length of the local regions.

As the local haplotype reconstruction complexity scales linearly with the length of the local regions, we have developed the ”envp”-mode, which allows users to specify a frequency threshold. Nucleotide variations occurring less frequently than the defined threshold in the read data are automatically excluded from the analysis. Consequently, this approach effectively reduces the length of the local region under investigation (see Supplementary Material section S3 for further details).

### Local haplotype reconstruction

For each of the local alignments, VILOCA undertakes haplotype reconstruction, which is the core functionality of the tool. Local haplotype inference is based on merging two key ideas. First, we exploit the information in co-occurring mutations by considering the full read sequence (detailed below in the section Mathematical model), rather than each sequence position separately. Second, we use the sequencing quality scores to assess how likely it is that an observed nucleotide variation is a true mutation as opposed to a technical sequencing error.

The Phred-scaled quality score *Q* represents the confidence that the sequencer called the base correctly. It is related to the error probability *E* as *Q* = *−*10 log *E*. The probability that a *_–_ <u>Q</u>* base is called correctly is then given by 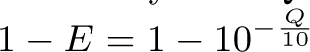 [21]. If Phred-scaled quality scores are not available, we instead learn a uniform error rate from the data (Supplementary Material section S2).

Since the number of haplotypes present in the sample is unknown, we utilize a Dirichlet process mixture model for local haplotype reconstruction. This model has the property that the number of haplotypes can be learned from the data. It assumes that the observations originate from a mixture of distributions, enabling the clustering of observed error-prone reads around their unobserved haplotypes. Cluster centers correspond to the unobserved haplotypes and the number of reads assigned to each cluster is proportional to the relative frequency of the corresponding haplotype in the population. Sequencing errors are identified as the discrepancies between bases in reads and their assigned haplotypes.

#### Mathematical model

After read alignment we can assume a local region of length *L* base pairs and *N* aligned reads *r* = (*r*_1_*, . . . , r_N_*) constraint to the region, such that *r_n_* = (*r_n,_*_1_*, . . . , r_n,L_*), for all *n* = 1*, . . . , N* . The per-position error rates *E* = (*E*_1_*, . . . , E_N_*) with *E_n_* = (*E_n,_*_1_*, . . . , E_n,L_*), for all reads *n* = 1*, . . . , N* , are obtained from the Phred-scaled quality scores as described above. There is no quality score for a deletion, therefore we assign to it the quality score of its 5’ neighboring nucleotide.

A Dirichlet Process Mixture Model allows an infinite number of clusters in the mixture. Instead of utilizing the complete Dirichlet process mixture model, we opt for the truncated version, which restricts the number of clusters, resulting a finite model size. This truncated approach serves as an effective approximation of the full model, particularly when a sufficiently large maximum number of clusters is selected [22]. For this purpose, we introduce the maximal number of haplotypes *K* and consider the mixing proportions 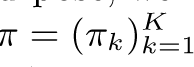 which are the relative haplotype frequencies. Let *z* = (*z*_1_*, …, z_N_*) be the haplotype assignments of the reads: if *z_n_* = *k*, then read *n* is obtained from haplotype *k*. We also write *z_nk_* = 1 in this case, i.e., if read *n* is assigned to cluster *k*, and zero otherwise. We place a symmetric Dirichlet prior on the mixing proportions 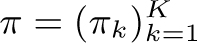

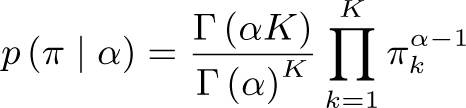

with concentration parameter *α*. The probability of the cluster assignment of read *n* is then

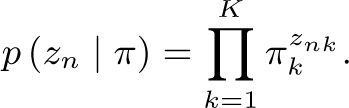

The *K* haplotypes are denoted by *h* = (*h*_1_*, …, h_K_*), where *h_k_* = (*h_k_*_1_*, …, h_kL_*), for all *k* = 1*, . . . , K*. As we are handling aligned DNA reads the alphabet is given by *B* = *{A, C, G, T, −}* and *| B |*= 5 is the size of the alphabet. In the following, it is convenient to encode the reads and haplotypes by indicator vectors: We write *r_nlb_*= 1 if alphabet letter *b* is present at position *l* in read *n*, and zero otherwise. And we write *h_klb_* = 1 if letter *b* is at position *l* in haplotype *k*, and zero otherwise.

To place a prior on the haplotype sequences, we assume that they are generated by mutation from a master sequence, denoted *h*_0_. Specifically, haplotypes are generated by introducing mutations into *h*_0_ with the per-position mutation rate 1 *− γ*, such that

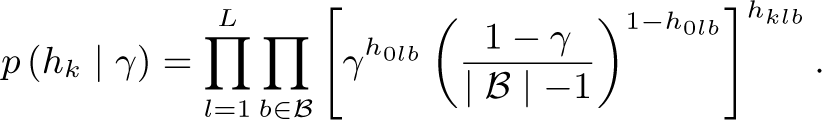

The mutation parameter *γ* is drawn from a Beta distribution Beta(*t*_0_*, t*_1_) with the hyperparameters *t*_0_ and *t*_1_.

The observed sequencing reads are then generated as erroneous copies of the haplotypes. The error process is governed by the position- and read-specific error probabilities *E* = (*E_n,l_*). The probability of observing a read *r_n_* is

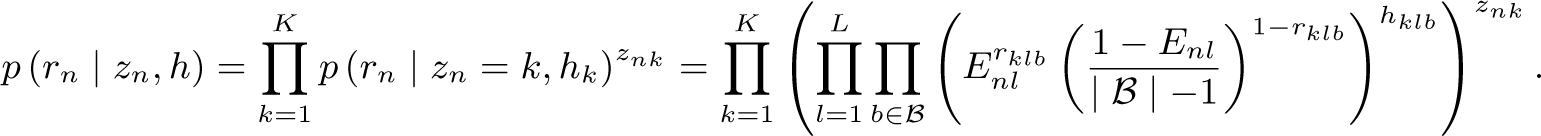

#### Variational inference

We aim to find the posterior probability distributions of the unobserved variables in the model, in particular of the haplotype sequences and frequencies. One approach would be to iteratively draw samples which will ultimately give us the shape of the joint posterior distribution of the unknown variables. However, this approach is usually slow and often requires prohibitively many samples to characterize the distribution correctly. Instead, we employ a variational inference approach. We developed a mean field approximation for the posterior which provides an analytical solution for the approximation. We assume that the latent variables are independent of each other, and each variable is described by its own variational distribution,

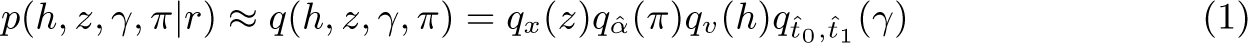

where *x*, *v*, *α*^, *t*^^^_0_, and *t*^^^_1_ are the parameters describing the variational distributions *q*. To find the best approximation, we minimize the Kullback-Leibler divergence between the posterior and the variational distribution, which is equivalent to maximizing the evidence lower bound (ELBO; see details in Supplementary Material section S1),

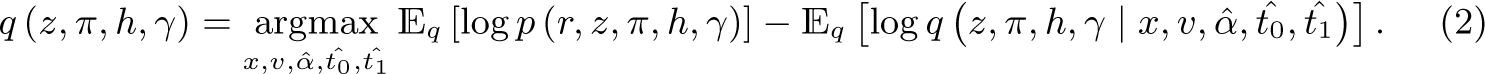

In order to solve this optimization problem, we have developed a coordinate ascent variational inference algorithm (Supplementary Figure S1), where the parameters defining the approximate distributions are iteratively learned from the data (Supplementary Material section S1). For each iteration the computational complexity is given by *O*(*NL*) and is hence scaling linearly with the coverage and the length of the local region. Once the algorithm has reached convergence, it outputs the predicted haplotypes along with their estimated relative frequencies. Additionally, it provides the posterior probability of each haplotype, which serves as a confidence score.

### Post-processing with strand bias filter

After local haplotypes have been predicted for each local region, mutation calls are generated by comparing the reconstructed haplotypes to the reference sequence. Each mutation call is assigned the posterior probability of the corresponding haplotype on which it occurred. For Illumina paired-end reads, the strand bias test developed by McElroy et al. [23] is applied to the mutation calls. The test assumes that mutations that are significantly more abundant on forward compared to reverse reads, or vice versa, are likely sequencing errors. Those variations are filtered out. Only mutation calls that pass both the strand bias test and a user-defined posterior probability threshold are reported.

### Benchmarking study

To assess the performance of VILOCA, we extended and applied the benchmarking module of V-pipe 3.0 [20]. The module facilitates the assessment of viral diversity estimation techniques from NGS data through flexible options for simulating ground truth haplotype populations and resulting sequencing reads, and for metrics to compare performance. To apply the module in our specific setting we integrated the read simulator PBSIM2 [24], which enables the simulation of sequencing read data from the Oxford Nanopore and PacBio platforms. Furthermore, we incorporated the tool primalscheme [25] to facilitate the simulation of Illumina sequencing reads using primer schemes.

#### Simulated datasets

We generated 10 samples of simulated Illumina sequencing reads with two different primer schemes to illustrate different VILOCA tiling strategies. We extracted a 5000bp region from the HIV HXB2 strain as reference sequence and simulated 15 haplotypes with pairwise distances between 4 *−* 12% and mutation frequencies between 0.5 − 25%. We set the amplicon size to 400bp and the amplicon overlap size to 10 and 100bp, referred to as scheme A and B, respectively.

Additionally, we generated a dataset simulating 60 different sequencing samples and comprising Illumina, PacBio, and ONT sequencing reads at varying coverage and complexity levels (Table 1). As reference sequence for the haplotype populations, we extracted a 249bp region for Illumina samples and a 5000bp region for ONT and PacBio samples from the HIV-HXB2 strain. These choices allow us to evaluate the performance of VILOCA directly in terms of local haplotype reconstruction for the respective sequencing technologies.

**Table 1:**
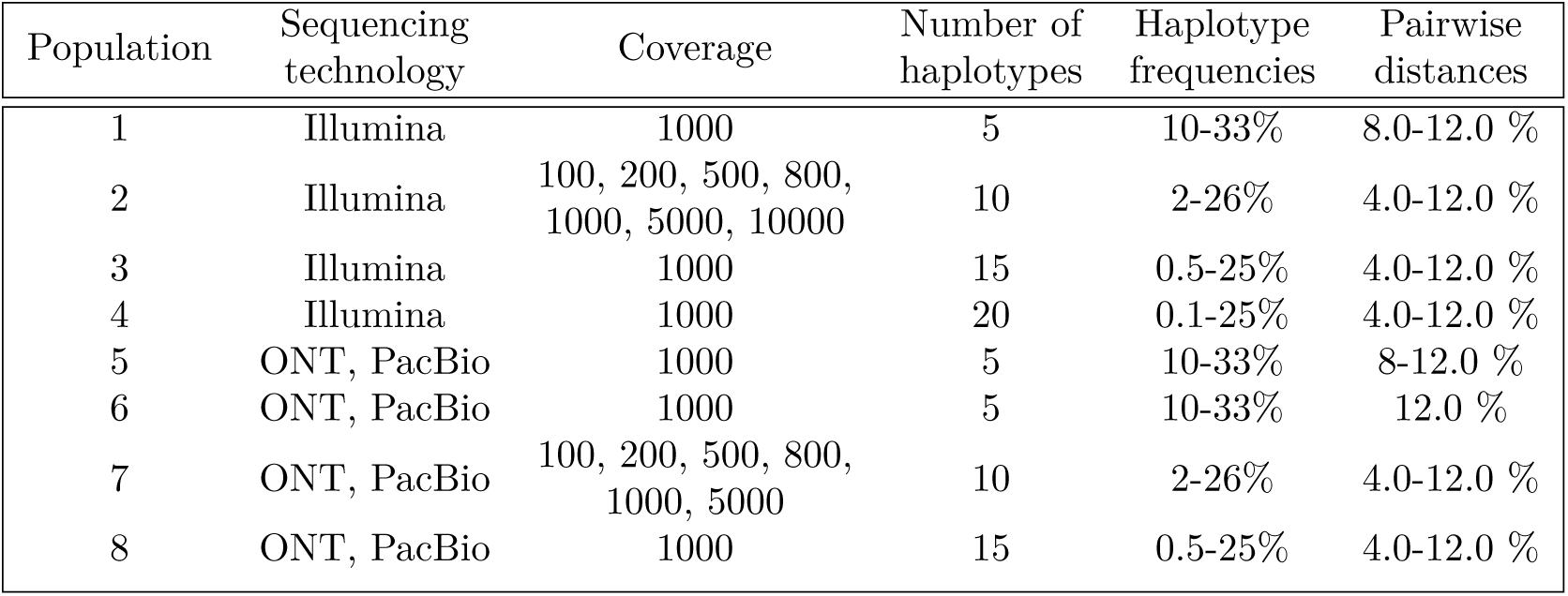
Simulated datasets with their population structure. We generated 5 replicates for each haplotype population configuration. Extended version of this table is the the Supplementary Material Table S1. Datasets were simulated using V-pipe’s benchmarking module [20].

| Population | Sequencing technology | Coverage | Number of haplotypes | Haplotype frequencies | Pairwise distances |
| --- | --- | --- | --- | --- | --- |
| 1 | Illumina | 1000 | 5 | 10-33% | 8.0-12.0 % |
| 2 | Illumina | 100, 200, 500, 800,<br>1000, 5000, 10000 | 10 | 2-26% | 4.0-12.0 % |
| 3 | Illumina | 1000 | 15 | 0.5-25% | 4.0-12.0 % |
| 4 | Illumina | 1000 | 20 | 0.1-25% | 4.0-12.0 % |
| 5 | ONT, PacBio | 1000 | 5 | 10-33% | 8-12.0 % |
| 6 | ONT, PacBio | 1000 | 5 | 10-33% | 12.0 % |
| 7 | ONT, PacBio | 100, 200, 500, 800,<br>1000, 5000 | 10 | 2-26% | 4.0-12.0 % |
| 8 | ONT, PacBio | 1000 | 15 | 0.5-25% | 4.0-12.0 % |

To generate the haplotype populations, we introduced variation by altering pairwise distance between haplotypes and the number of haplotypes, leading to diverse haplotype frequency spectra (Table 1). The frequencies within each spectrum correspond to the terms of the geometric series starting with the ratio of 0.75, resulting in the presence of a small number of high-frequency haplotypes and a larger number of low-frequency haplotypes as predicted by viral quasispecies theory [26]. For each population configuration, we generated five haplotype population replicates. This strategy resulted in populations with 5 − 20 haplotypes at frequencies between 0.1% and 33%.

#### Real data

We used three experimental datasets in our benchmarking study (Table 2). The Five-HIV-Mix dataset consists of Illumina MiSeq sequencing data derived from a mixture of five HIV-1 strains: HXB2, 89.6, JR-CSF, NL4-3, and YU-2 with approximately equal frequencies of 20% [27].

**Table 2:**
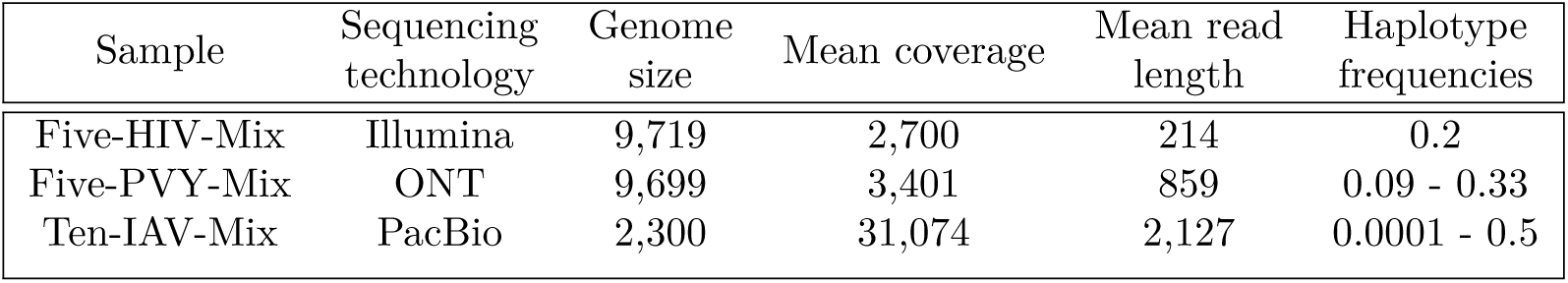
Overview of the four experimental samples used in the benchmarking study.

| Sample | Sequencing technology | Genome size | Mean coverage | Mean read length | Haplotype frequencies |
| --- | --- | --- | --- | --- | --- |
| Five-HIV-Mix | Illumina | 9,719 | 2,700 | 214 | 0.2 |
| Five-PVY-Mix | ONT | 9,699 | 3,401 | 859 | 0.09 - 0.33 |
| Ten-IAV-Mix | PacBio | 2,300 | 31,074 | 2,127 | 0.0001 - 0.5 |

The Five-PVY-Mix dataset comprises an artificial mixture of ONT MinION sequencing samples (accession numbers: SRR11431597, SRR11431596, SRR11431617, SRR11431616, SRR11431615) from five Potato virus Y strains (GenBank accession numbers MT264731–MT264735) [28]. Frequencies of the strains range from 9% to 33%. To define ground truth mutations, we compared these strains to our reference strain *NTNa* (GenBank accession number: MT264732). The limited read length in this dataset covers only approximately 9% of the full genome, allowing for SNV performance assessment but rendering it unsuitable for evaluating haplotype reconstruction (Table 2).

Finally, the Ten-IAV-Mix dataset is a PacBio sample containing ten Influenza A virus strains [29]. The strains have frequencies ranging from 0.1% to 50% [16]. Since a significant fraction of the reads span the entire genome, we were able to run VILOCA with a single local region covering the full genome, effectively allowing us to evaluate global haplotype reconstruction.

### Comparison to existing methods

VILOCA provides mutation calls and local haplotypes for Illumina, ONT, and PacBio sequencing data. To ensure a comprehensive evaluation and meaningful comparison against state-of-the-art methods, we included relevant tools in our study that provide those outputs.

LoFreq (version 2.1.5, [30]) is a widely used mutation caller compatible with all three sequencing technologies. Similarly to VILOCA, LoFreq incorporates sequencing quality scores in its error model. However, unlike VILOCA, it is restricted to single base-pair mutation calls and does not support local haplotype inference. LoFreq was executed with default settings.

ShoRAH (version 1.99.2, [17]) provides mutation calls and local haplotypes. However, ShoRAH is primarily optimized for Illumina reads, lacks flexibility in tiling the genome into local regions, and does not consider sequencing quality scores to inform its intrinsic error model. ShoRAH was executed with default settings.

For comparison with our haplotype predictions, we also considered global haplotype reconstruction methods. CliqueSNV (version 2.0.2, [16]) is among the best performing haplotype callers in recent benchmarking studies [19, 20] and specialized for Illumina and PacBio sequencing samples. We executed CliqueSNV in *snv-illumina* and *snv-pacbio* mode. The original publication claims that it also obtains reliable results on ONT samples, which we also assessed in this study. If not stated differently, we executed CliqueSNV setting the parameter *tf* to 0.001 (see details in Supplementary Material section S4.2). This parameter is the minimal threshold for haplotype frequencies, haplotypes or mutation with frequency smaller than this threshold will not be detected.

Additionally, we included the single-end read version of PredictHaplo (version 1.1, [18]) to reconstruct haplotypes from the long-read sequencing samples.

Finally, we attempted to evaluate the non-reference guided global haplotype reconstruction method Strainline [15]. However, we encountered an error in the clustering step that prevented us from running Strainline successfully (https://github.com/HaploKit/Strainline/issues/ 17).

### Performance measures

We assessed the performance of all methods in terms of their ability to (a) call single-nucleotide variants (SNVs), and (b) reconstruct local haplotype sequences, along with the accuracy of their estimated relative frequencies in the sample. For SNVs, precision, recall, and f1 scores are computed by comparing the predicted with the true nucleotide. Local haplotype reconstruction is evaluated by calculating precision and recall for the recovered ground truth haplotypes, considering only the corresponding local regions. To determine the accuracy of predicted haplotypes, we compared them to the ground truth haplotypes and measured the Hamming distance. If the Hamming distance (normalized by sequence length) is below a certain threshold value *λ*, the predicted haplotype is considered correct, i.e., a true positive; otherwise, it is marked as a false positive. False negative haplotypes are ground truth haplotypes that do not match any predicted haplotype (with a Hamming distance smaller than *λ*). We set the threshold *λ* to 0.01, thus tolerating up to 1% of nucleotides to deviate for a haplotype match.

## 3 Results

We first present the results of the benchmarking study, and in the second part, we demonstrate applications of VILOCA in three different real-life scenarios.

### 3.1 Benchmarking study

We evaluated the performance of different VILOCA variations on the set of 10 samples with varying primer schemes for the Illumina read simulation. First, we compared the performance of VILOCA in envp mode with different thresholds. In this mode, positions with nucleotide variations occurring less frequently than the defined threshold are excluded from the analysis. We noticed that executing VILOCA in this mode with thresholds of 0.01 and 0.001 does not change performance significantly (Figures 2A). However, if users are not interested in mutations below a certain frequency threshold, running VILOCA with an envp threshold can reduce runtime drastically (Figures 2B). Next, we varied the primer scheme for Illumina read simulation between the 10 datasets. In terms of precision, there was not much difference between the two schemes. However, for recall, there were performance differences depending on the tiling strategy used. If there is a rather small overlap between amplicons it is beneficial if users can provide the primer scheme to VILOCA as input (Scheme A, Figure 2A). Contrarily, when reads are spread more uniformly along the genome it is beneficial in terms of recall to apply the uniform tiling strategy (Scheme B, Figure 2A).

**Figure 2:**
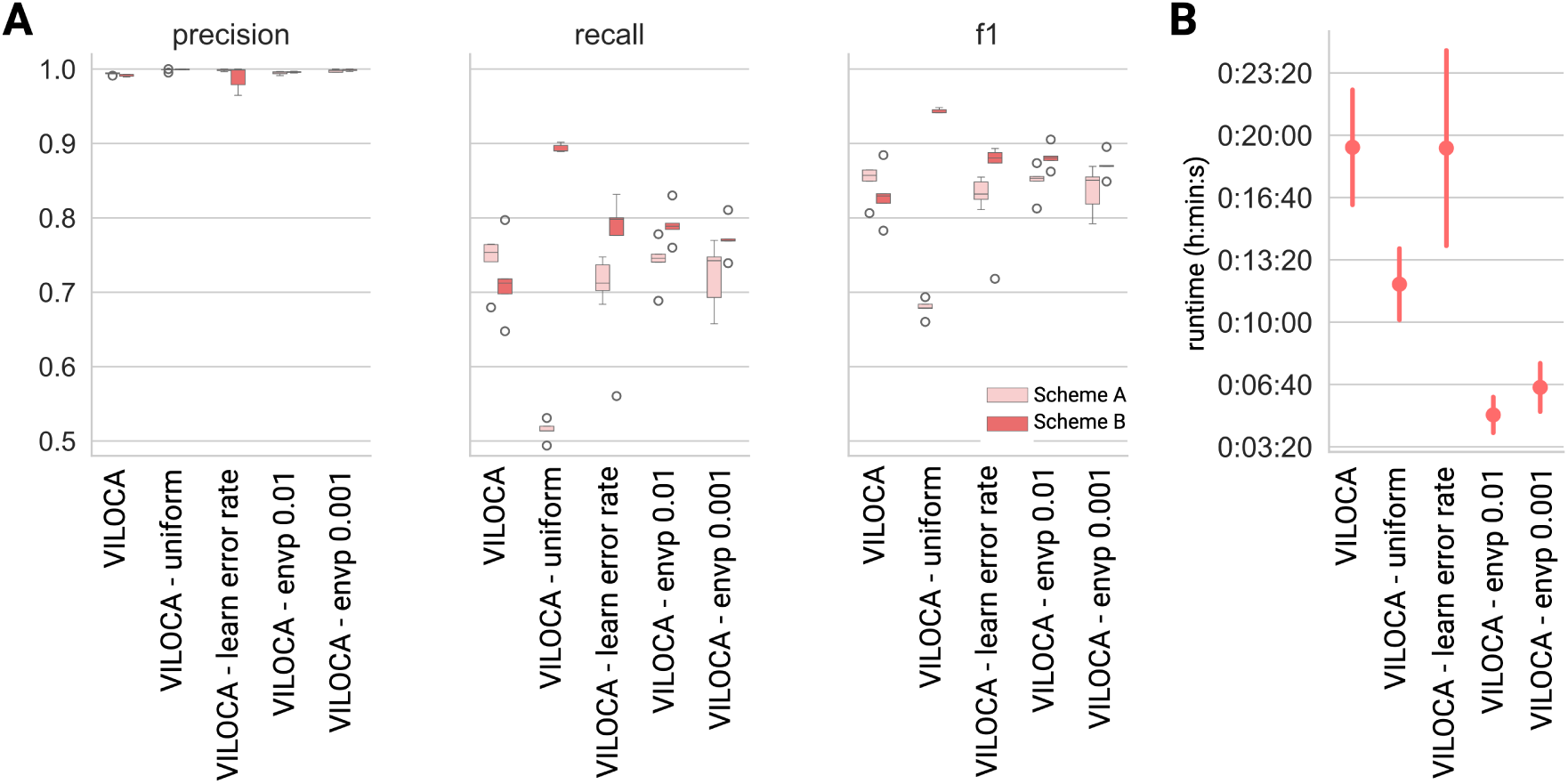
Comparison of VILOCA executed in different modes: VILOCA (primer tiling strategy), VILOCA - uniform (uniform genome tiling strategy), VILOCA - learn error rate (primer tiling strategy with error rate learned from the data), and VILOCA - envp (primer tiling strategy with envp-mode, thresholds of 0.01 and 0.001). In envp-mode, positions with nucleotide variations occurring less frequently than the defined threshold are excluded from the analysis. The VILOCA modes were evaluated on 10 samples of simulated Illumina sequencing reads with varying primer schemes read simulation. (A) Precision, recall, and f1-score comparison of the different VILOCA modes with respect to the different primer schemes applied. Scheme A (pink) represents a primer scheme with an amplicon size of 400 bps and an overlap of 10 bps. Scheme B (red) has an overlap of 100 bps. (B) Runtime comparison of the different VILOCA modes.

Since VILOCA is providing a posterior probability for each mutation call, we evaluated the effect of varying the thresholds of the posterior on the performance on the simulated short- and long-read datasets described in Table 1. A low posterior threshold will increase recall, whereas a stricter, i.e. higher, threshold will increase precision. This trade-off is indeed observed when varying the posterior threshold (Supplementary Figure S3 and S4). Our analysis suggests setting the posterior thresholds to 0.9 for Illumina samples, and to 0.5 for the long-read sequencing samples (Supplementary Figure S3 and S4).

#### Mutation calling performance on simulated datasets

We first analysed mutation calls in terms of precision, recall and f1-score. Across all simulated datasets, we found that VILOCA is performing best for the Illumina samples, whereas there is more variation for the long-read samples (Figure 3A). This can be explained by the approximately one magnitude lower error rate compared to the long-read sequencing technologies. In our simulated long-read datasets the sequencing error rate is around 0.15; in contrast, the Illumina samples have an error rate around 0.001. Both LoFreq and VILOCA exhibit higher precision than recall. In long-read samples, VILOCA demonstrates significantly higher recall compared to LoFreq which can be attributed to their different approaches (Figure 3A). LoFreq treats each sequence position independently, making it impossible to leverage additional information from co-occurring mutations. By contrast, both ShoRAH and VILOCA consider entire reads, enabling them to exploit the additional information provided by co-occurring mutations. Since PacBio and ONT samples contain longer reads, which inherently carry more of such information, these tools are able to mitigate the noisier nature of long-read data. When comparing the estimated mutation frequencies with the ground truth values, we observe that VILOCA provides highly accurate predictions for both short- and long-read data (Figure 3A, bottom row). To analyse the effect coverage on the performance, we simulated sequencing reads for Population 2 and 6 with varying coverage. The performance gain of VILOCA is particularly pronounced for low-coverage samples, and shows a general trend of improved recall as coverage increases (Figure 3B).

**Figure 3:**
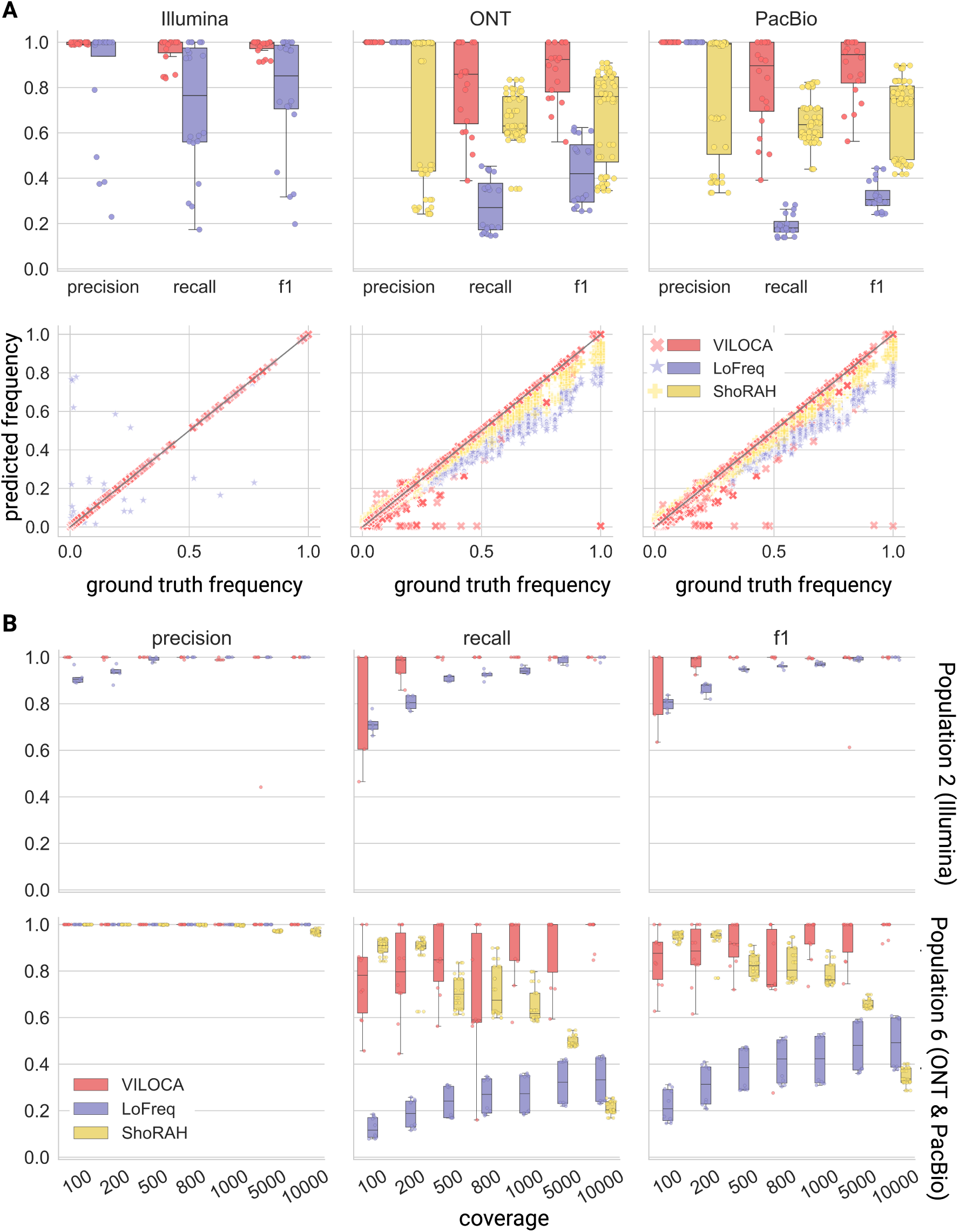
Performance evaluation in terms of mutation calling for simulated long and short reads. (A) Top row: Precision, recall and f1-score for mutation calls on simulated Illumina, ONT, and PacBio datasets with coverage 1000. Bottom row: Comparison of mutation frequency predictions to ground truth for simulated Illumina, ONT, and PacBio datasets as specified in Table S1. Each data point represents a true positive mutation call. The black line denotes perfect alignment between predicted and ground truth frequencies. (B) Mutation calling performance at varying coverages of 100, 200, 500, 800, 1000, 5000, and 10000bp for Population 2 (short-read samples; top row) and Population 6 (long-read samples; bottom row).

#### Haplotype reconstruction on simulated datasets

For haplotype reconstruction, in accordance with its mutation calling performance, VILOCA is exhibiting generally higher precision and recall for the short-read samples compared to the long-read samples. For short and long-read samples, VILOCA has near perfect precision remaining above 0.9 for all populations (Figure 4A+B). The recall of VILOCA is generally decreasing with increasing number of haplotypes in the population (Figure 4A+B). Especially for the long-read samples, the recall is decreasing drastically to 0.42 for Population 8 (Figure 4B). The decrease can be ascribed to the composition of the population, which includes 3 haplotypes below 1% frequency and 8 haplotypes with frequencies below 5%. With the noise present in long-read sequencing data, detecting these low-frequency haplotypes becomes challenging. This can be also observed for the predictions of PredictHaplo. For Population 7 and 8, PredictHaplo is rather detecting consensus sequences of the each haplotype cloud, and is not able distinguish the single haplotypes (Population 7 and 8, Figure 4B). CliqueSNV failed in achieving convergence for the datasets of simulated PacBio reads and only reported consensus sequences for the samples. For further details, refer to the Supplementary Material section S4. For the ONT samples, the performance is very similar with no significant differences (Supplementary Figure S5).

**Figure 4:**
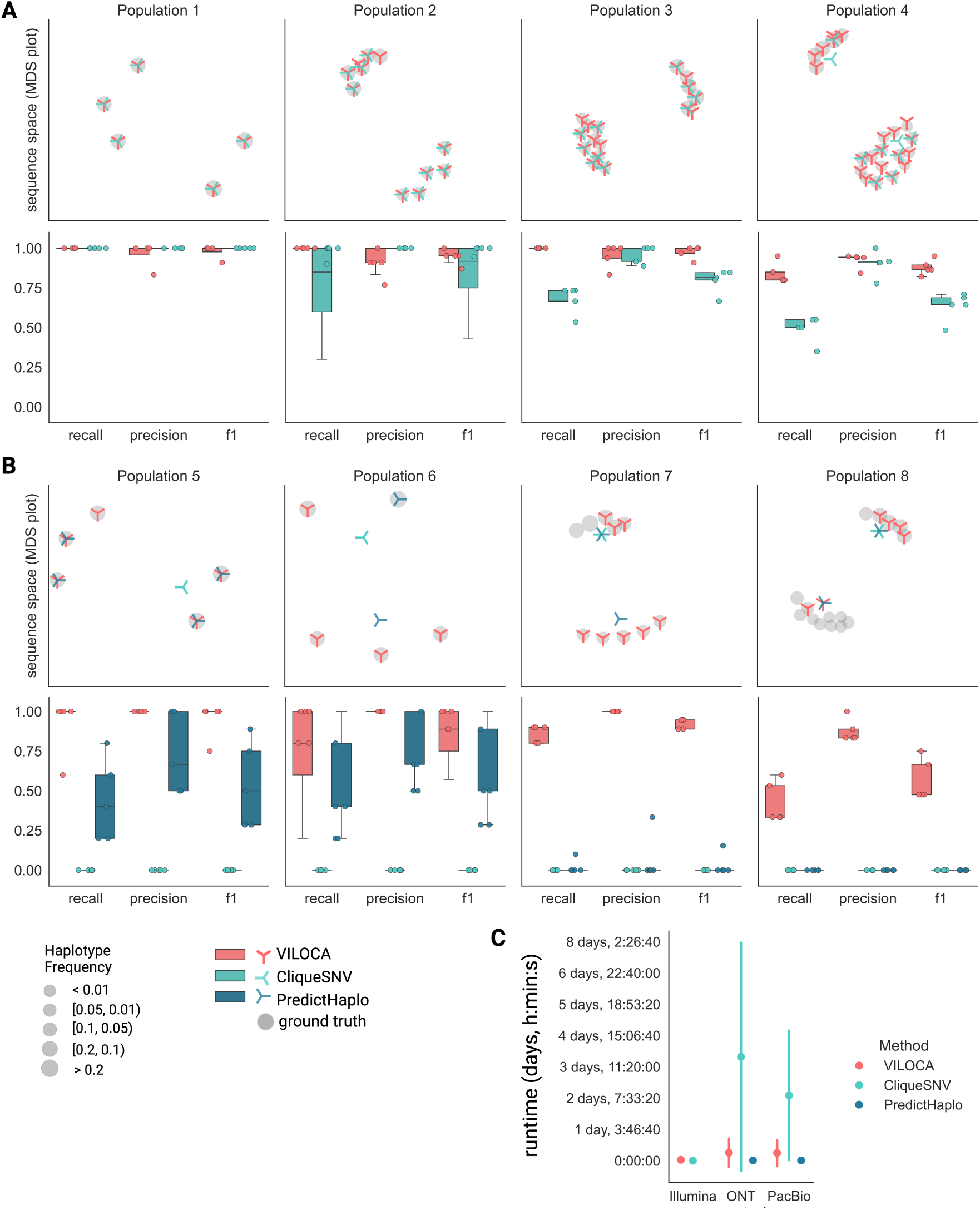
Haplotype reconstruction performance on datasets of simulated Illumina (A) and PacBio (B) sequencing reads. Top row: Multidimensional scaling (MDS) plots for the sequence space visualization of one example simulation replicate per haplotype population. Each marker represents one haplotype sequence and marker size corresponds to haplotype frequency in the sample. Bottom row: Precision, recall and f1-score of haplotype inference across the five replicates. For the computation of true and false positives, we employ a distance threshold of 0.01 between predicted and ground truth sequence. (C) Runtime comparison of the three methods across all simulated datasets of Illumina, ONT, and PacBio reads. Error bars are depicting the standard deviation.

#### Runtime comparison

To compare the runtime between different methods on the simulated datasets, we executed them allocating equivalent computational resources through our Snakemake benchmarking workflow. The maximum runtime for VILOCA was 2 days and 18 hours when analyzing long-read samples with local regions of 5000 base pairs and a coverage of 10000 reads, while PredictHaplo had a maximal runtime of less than 10 hours (Figure 4C). For the long-read samples, CliqueSNV demonstrates the longest runtime, which can be attributed to its quadratic relationship with the number of SNVs per read, as discussed in Supplementary Material section S4. Between the tools reporting mutations, LoFreq demonstrated the shortest runtime, which is expected as positions are treated independently and the method is therewith highly parallelizable (Supplementary Figure S6).

#### Performance on real experimental samples

We applied all methods to the three experimental datasets (Table 2), for which the underlying viral strains are known. All three datasets were used for the evaluation of mutation calling. Given that the Ten-IAV-Mix comprises a significant proportion of reads spanning the entire genome, it enabled also the assessment of the haplotype predictions of VILOCA. Across all datasets, VILOCA excels in mutation calling with precision values exceeding 0.9 (Figure 5 A). VILOCA outperforms other methods in the Five-PVY-Mix and Five-HIV-Mix datasets in terms of f1-score. In the Ten-IAV-Mix dataset, both LoFreq and VILOCA exhibit high precision above 0.95 but low recall under 0.5 (Figure 5 A). The low recall for this sample can be attributed to the increased number of haplotypes with frequencies below 1% (Table 2). ShoRAH fails to provide meaningful predictions for this dataset, yielding 0 precision and recall. In terms of haplotype reconstruction, VILOCA identifies five out of ten haplotypes in the Ten-IAV-Mix without predicting any false positives (Table 3, Figure 5 B). The undetected haplotypes have frequencies below 0.05, and have not been detected possibly due to VILOCA excluding approximately 10% of reads from the analysis, as they do not cover a big enough fraction of the genome.

**Figure 5:**
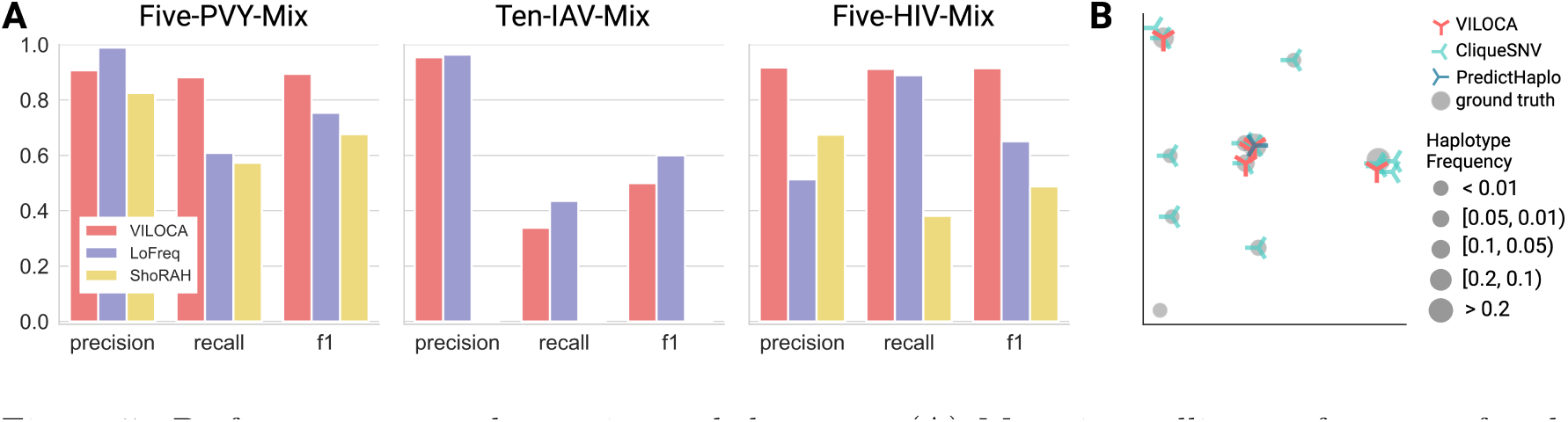
Performance on real experimental datasets. (A) Mutation calling performance for the Five-PVY-Mix (ONT), the Ten-IAV-Mix (PacBio), and the Five-HIV-Mix (Illumina), for VILOCA, LoFreq, and ShoRAH. (B) MDS visualization of haplotypes in sequence space. Each mark represents a haplotype. Symbol size corresponds to the frequency of the respective haplotype in the sample. CliqueSNV was executed with setting the parameter *-tf* to 0.001 to allow detection of low-frequency haplotypes.

**Figure 6:**
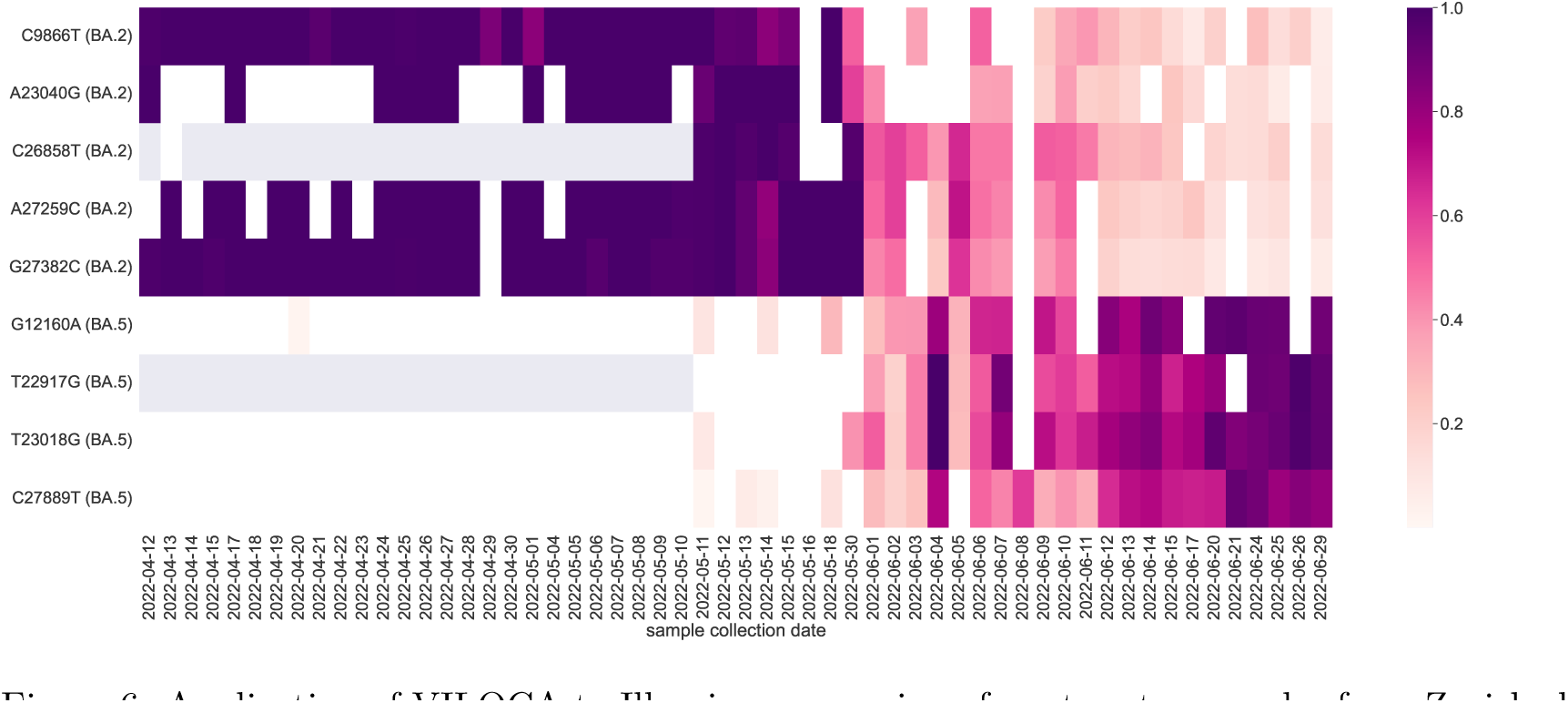
Application of VILOCA to Illumina sequencing of wastewater samples from Zurich shows the emergence of Omicron BA.5 variant while the BA.2 variant is accordingly decreasing. The heatmap shows the mutation frequencies provided by VILOCA for each sample. We only include the mutations that are specific to either BA.2 or BA.5, no shared mutations are reported in this analysis. Gray cells mark that no reads covered the respective position in the sample. The first detection of a BA.5-specific mutation is on 2022-04-20.

**Table 3:**
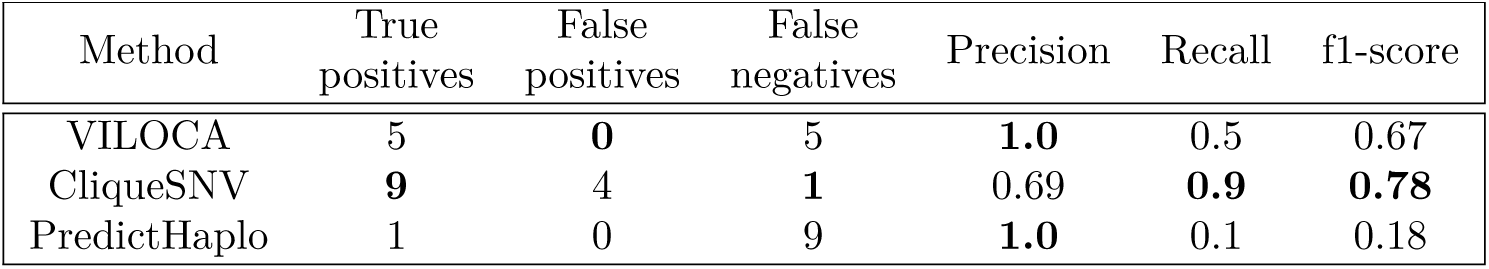
Haplotype reconstruction performance on the Ten-IAV-Mix dataset. With distance threshold of 0.001 corresponding to less than 3 mismatches in predictions compared to the ground truth.

| Method | True positives | False positives | False negatives | Precision | Recall | f1-score |
| --- | --- | --- | --- | --- | --- | --- |
| VILOCA | 5 | **0** | 5 | **1.0** | 0.5 | 0.67 |
| CliqueSNV | **9** | 4 | **1** | 0.69 | **0.9** | **0.78** |
| PredictHaplo | 1 | 0 | 9 | **1.0** | 0.1 | 0.18 |

### Applications

To demonstrate the versatile uses of VILOCA, we applied it to three different real-world datasets. The data processing details are in Supplementary Material section S5.

#### Early detection of low-frequency mutations in patient samples

To demonstrate application to clinical samples, we analysed samples from an HIV-1-positive patient, published in [31]. The patient was part of the CAPRISA 002 cohort in South Africa, and several longitudinal samples pretreatment are available. We analysed six longitudinal pre-treatment Illumina samples from week 30 to 240 post-infection (Supplementary Table S2). Using VILOCA, we were able to call mutations at frequencies below 1%, which subsequently increase in later samples. For instance, VILOCA detects mutation C982T at 0.5% frequency with high posterior probability of 0.96 at week 30 post-infection, and the mutation rises to 15% frequency with posterior of 1.0 by week 240, supporting its earlier presence in the population (Supplementary Figure S7). By contrast, LoFreq only detects the mutation at week 135 post-infection (Supplementary Figure S7). The ability of VILOCA to identify mutations at low frequencies and hence early in the course of evolution is important, for example, for mutations associated with drug resistance or immune escape.

#### Detection of a newly emerging SARS-CoV-2 variant in sewage samples

To showcase that VILOCA can also handle settings of very high noise, we applied it to data obtained from wastewater samples produced in the scope of the Swiss SARS-CoV-2 Wastewater Surveillance [32]. To detect newly emerging SARS CoV-2 variants, we processed 56 wastewater samples that were collected in the period of 2022-04-12 until 2022-06-26 in Zurich and sequenced using Illumina technology following the ARTIC protocol [25] (ENA Project accession number: PRJEB44932). Applying VILOCA we observed the emergence of the SARS-CoV-2 variant BA.5 and the fading of the BA.2 variant. Specifically, on 2022-04-20, we called the first BA.5-specific variant mutation G12160A at 1.5% frequency and with perfect posterior probability of 1.0 (Figure 6) indicating highest confidence in the mutation call. Subsequently, this mutations occurs again on 2022-05-11 at 10% frequency and together with two other BA.5-specific mutations (T23018G and C27889T) with frequencies 8.5% and 1.5%, respectively, both with posterior 1.0. Concurrently, we noticed a continuous decrease in the BA.2-specific mutations C9866T, A23040G, C26858T, A27259C, and G27382C. In comparison, COJAC [5] the method employed by CovSpectrum [33] on that same data, is predicting the first occurrence of BA.5 SARS-CoV-2 variant in Switzerland to be on 2022-05-11, i.e., almost 3 weeks later. Unlike VILOCA, COJAC relies solely on the mutation occurrence in the raw reads and does not provide error correction and confidence scores. Hence, VILOCA improves early detection of newly emerging variants in noisy wastewater sequencing data.

#### Identification of circulating polio strains in a sewage sample

To showcase VILOCA an application on long-read sequencing data, we demonstrate the detection of mixtures of Poliovirus strains and Coxsackievirus from MinION sequencing sewage samples. We analysed a sewage sample (accession number: ERR4033236) from Pakistan that was collected in the scope of the official polio surveillance program by the WHO and Pakistan Government [8]. To differentiate between various polio types, Oxford Nanopore MinION sequencing targeting the VP1 region was employed [8]. VILOCA was applied to a single local window of the genome spanning the entire VP1 region as most reads spanned vast parts of the VP1 region. VILOCA predicted five high-confidence haplotypes covering the full VP1 region (posterior probability *≥* 0.95). Using BLAST [34] we mapped the predicted haplotypes and identified a mixture of wild poliovirus type 1, Sabin poliovirus type 2, and Coxsackievirus A13 (Table 4). These findings align with the results presented by Shaw et al. [8]. The authors also identified the presence of Coxsackievirus A13 and Poliovirus 2 in this and other sewage samples. Further, in Pakistan, a total of 54 cases of wild poliovirus type 1 were reported in 2015, respectively [35], and other studies also confirm its detection in environmental samples [35]. The identification of the Sabin vaccine virus is consistent with the administration of oral poliovirus vaccine routines during the same period from 2015 in vaccination campaigns across Pakistan [36, 35, 37]. Thus, our findings highlight that VILOCA can be applied also to MinION sequencing data, and near full-length haplotypes can be generated for surveillance and detection of circulating viral strains.

**Table 4:** Identification of circulating Poliovirus and Coxsackievirus in a sewage sample from Pakistan. VILOCA reconstructed five high-confidence haplotypes of the VP1 region of the genome with posterior probability *≥* 0.95. Using BLAST [34] the predicted haplotypes were mapped against the Nucleotide collection database.

| Haplotype ID | No. of reads | BLAST hit | Identity |
| --- | --- | --- | --- |
| 1 | 281 | Poliovirus 2 strain Sabin-like [42] (MG808048.1) | 96.04% |
| 2 | 323 | Poliovirus 2 strain Sabin (AY184220.1) | 95.09% |
| 3 | 178 | Poliovirus 2 strain Sabin (AY184220.1) | 96.27% |
| 4 | 553 | Coxsackievirus A13 (JF260922.1) | 81.27% |
| 5 | 456 | Poliovirus 1 (AY082690.1) | 83.59% |

## 4 Discussion

The estimation of viral genetic diversity is crucial in understanding viral dynamics, disease patterns, and treatment outcomes. Previous studies have primarily focused on SNVs to assess viral diversity. However, analyzing co-occurring mutations and local haplotypes offers a more comprehensive and informative approach. Unlike SNVs, which capture individual nucleotide changes, examining co-occurring mutations and local haplotypes provides insights into multiple mutations occurring (locally) on the same genetic background. This piece of information is particularly valuable when analyzing viral populations with complex epistatic interactions affecting viral phenotypes such as virulence [38], viral fitness [39], drug resistance [2, 3], and immune escape [40, 41].

Global haplotype reconstruction methods aim to infer full-length sequences by connecting reads based on their mutation profiles. While this approach provides a complete picture of variations present in the genome, the reconstruction is often computational expensive and can be challenging due to distant linked reads and stretches of homogeneity [20]. On the other hand, local haplotype reconstruction methods focus on smaller genomic regions, typically of length comparable to the read length. By limiting the scope to local regions, these methods can leverage the direct support of sequencing data to accurately reconstruct haplotypes within these regions with confidence. Additionally, utilizing information from co-occurring mutations though the reconstruction of local haplotypes enhances mutation calling performance results.

VILOCA focuses on generating local haplotypes. The selection of local region sizes depends on the available sequencing data. Typically, for Illumina sequencing data, one would choose the read length as the length of the region to be analyzed. However, if paired-end reads are available it is also an option to fuse paired-end reads into a single merged read and use the merged reads as input to VILOCA. This approach enables the reconstruction of extended regions. In contrast, selecting region lengths for long-read sequencing data is more complex due to significant variation in coverage and read length throughout the genome. Shorter regions yield higher coverage, while longer regions often have fewer supporting reads. Hence, the region length choice is always a tradeoff between higher coverage and longer regions, and the ultimate choice is depending on the specific research objectives. It might also be useful to run VILOCA with different region lengths on the same data.

In our benchmarking study, we have shown that VILOCA has high precision for local haplotype reconstruction on datasets of simulated Illumina reads while detecting haplotypes of *<* 1% frequency. VILOCA has demonstrated its capability to predict full-length haplotypes when long-read samples cover the entire genome. Even with increasing genetic diversity, VILOCA maintains high precision in its predictions using noisy long-read data. However, it encounters challenges in detecting low-frequency haplotypes in this setting. In terms of mutation calling, VILOCA outperformed the other methods, for both short-read and long-read samples. It accurately recovers mutations at low frequencies even on noisy long-read data by leveraging co-occurring mutations.

Compared to CliqueSNV, the runtime of VILOCA is determined by the coverage and size of the local regions rather than the diversity of the sample. This scalability allows VILOCA to process high-diversity long-read samples samples efficiently, while CliqueSNV fails on these samples. Nevertheless, the computational complexity can become a limitation when dealing with large amounts of long-read high-coverage samples. To address this, more scalable inference methods, such as Stochastic Variational Inference, for example, could be explored to improve runtime.

Furthermore, specific modeling decisions may be suboptimal in certain scenarios. For instance, the current model relies on sequencing quality scores as error probabilities, which may lead to inaccuracies if these scores are unreliable. While VILOCA offers the choice to adopt a uniform error rate and disregard sequencing quality scores, a more effective approach could involve learning position-specific error rates while still leveraging information from quality scores as a prior. Another potentially suboptimal modeling descision is the assumption of a single master sequence with a uniform mutation rate. This may be limiting in cases of highly diverse viral populations, like co-infections. To address this, the distribution over nucleotide bases per position as a prior could better account for sample heterogeneity.

To enhance user experience, we integrated VILOCA into V-pipe 3.0 [20], a computational pipeline tailored for analyzing viral genomes using next-generation sequencing data. This integration allows users to effortlessly input their raw reads into V-pipe 3.0 and retrieve mutation calls and local haplotypes produced by VILOCA as part of the pipeline’s output. By streamlining the analysis process, this integration offers users a comprehensive solution for studying viral genomes in an reproducible, scalable, and transparent fashion.

VILOCA finds practical applications in a wide range of fields, including classical mutation calling for single or longitudinal samples, and viral surveillance in wastewater. Its ability to accurately infer the structure of intra-host viral populations makes it a valuable tool for studying viral evolution and transmission dynamics and exploring the genome compositions of RNA viruses both within hosts and within mixed samples in environmental settings. With the ongoing advancements in long-read sequencing technologies, studying viral evolution is becoming increasingly feasible. In this evolving landscape, VILOCA provides improved accuracy in mutation and haplotype calling, particularly for long-read sequencing data.

## 5 Availability of Data and Code

The code base for VILOCA is publicly available on GitHub at https://github.com/cbg-ethz/ VILOCA. The benchmarking study, including all data-generating scripts, method execution, and notebooks for figure creation, can be accessed on GitHub: https://github.com/cbg-ethz/V-pipe/tree/feature-benchmark/resources/auxiliary_workflows/benchmark/resources/local_ haplotype_setup and can be reproduced by running the Snakemake workflow (Supplementary Material Section S4.1). The sequencing data of the Five-HIV-Mix is accessible on the Sequence Read Archive under the accession number SRX342666. The sequencing data of the Five-PVY-Mix is accessible on the Sequence Read Archive under the accession numbers SRR11431597, SRR11431596, SRR11431617, SRR11431616, SRR11431615. The sequencing data of the Ten-IAV-Mix is available on GitHub https://github.com/vtsyvina/CliqueSNV/ tree/master/data/PacBio_reads. The data processing pipelines used in the section Applications can be accessed on GitHub (https://github.com/cbg-ethz/SARS-CoV-2-wastewater-sample-processing-VILOCA/ tree/main/resources/setup_emergence_new_variant, https://github.com/cbg-ethz/viloca_applications). The repositories contain step-by-step instructions on how to reproduce the analysis and the presented figures. The accession numbers of the HIV Illumina samples used in the section Applications can be found in Supplementary Table S2. The SARS-CoV-2 samples analysed in the section Applications are under the ENA Project accession number PRJEB44932. A list of the sample accession numbers can be found in the Supplementary Table S3. The sewage sample from Pakistan can be found under accession number ERR4033236.

## Acknowledgements

Figures were created and assembled with BioRender (BioRender.com). LF was funded by European Union’s Horizon 2020 research and innovation program, under the Marie Skłodowska-Curie Actions Innovative Training Networks grant agreement No. 955974 (VIROINF).

