## Supplementary Material for "VILOCA: Sequencing quality-aware haplotype reconstruction and mutation calling for short- and long-read data"

Lara Fuhrmann[1,2], Benjamin Langer[1], Ivan Topolsky[1,2], Niko Beerenwinkel[1,2,*]

[1]Department of Biosystems Science and Engineering, ETH Zurich, Basel, 4058, Switzerland
[2]SIB Swiss Institute of Bioinformatics, Basel, 4058, Switzerland
*To whom correspondence should be addressed

#### S1 Model using the sequencing quality scores

In this section, we provide the detailed computations for the variational inference and the variational distributions that are used in our variational inference algorithm (Figure S1).

The aim of variational inference is to minimize the distance between the posterior distribution $p(h, z, \gamma, \pi \mid r)$ and the variational distribution $q(h, z, \gamma, \pi)$. We use the variational distribution given in Equation (1) in the main text which assumes independence of latent variables $z$, $\pi$, $h$, and $\gamma$, and hence gives rise to a mean field approximation. We solve the following optimization problem over the variational parameters

$$q(z, \pi, h, \gamma) = \underset{x,v,\hat{\alpha},\hat{t}_0,\hat{t}_1}{\operatorname{argmin}} \ \mathrm{KL}\left(q\left(h, z, \gamma, \pi \mid x, v, \hat{\alpha}, \hat{t}_0, \hat{t}_1\right) \parallel p\left(h, z, \gamma, \pi \mid r\right)\right).$$

The problem can be reformulated in terms of the evidence lower bound

$$\mathrm{ELBO}(q) = \mathbb{E}_{q(z,\pi,h,\gamma)}\left[\log p(z, \pi, h, \gamma, r)\right] - \mathbb{E}_{q(z,\pi,h,\gamma)}\left[\log q(z, \pi, h, \gamma)\right].$$

Maximizing the ELBO is equivalent to minimizing the KL-divergence:

$$\begin{aligned} q(Z) &= \operatorname{argmax} \mathrm{ELBO}\left(x, v, \hat{\alpha}, \hat{t}_0, \hat{t}_1\right) \\ &= \underset{x,v,\hat{\alpha},\hat{t}_0,\hat{t}_1}{\operatorname{argmax}} \ \mathbb{E}_q\left[\log p(r, z, \pi, h, \gamma)\right] - \mathbb{E}_q\left[\log q\left(z, \pi, h, \gamma \mid x, v, \hat{\alpha}, \hat{t}_0, \hat{t}_1\right)\right]. \end{aligned}$$

To solve this maximization problem, we derived a coordinate ascent variational inference algorithm (Figure S1). Through iterative updating, the parameters characterizing the variational distributions are learned. In each iteration, the expected value of each factorized variational distribution is updated while the remaining once are fixed. These update functions are derived using the general formula, as previously described [1],

$$\log q_i(Z_i) = \mathbb{E}_{m|m\neq j}\left[\log p(Z, r)\right] + \text{constant} \tag{1}$$

for each of the latent variables $Z = (h, z, \gamma, \pi)$ of the model. All terms not involving $Z_i$ are integrated into the constant terms as they are not changing with respect to $Z_i$. The algorithm guarantees that the ELBO is non-decreasing and converges to a local optimum. We measure the convergence by monitoring the absolute changes in the ELBO. In the following we derive the update functions for each distribution and the ELBO.

Using the general formula Equation 1, we find for the variational distribution of the cluster assignments:

$$
\begin{aligned}
\log q^*(z) &= \mathbb{E}_{q(h,\gamma,\pi)}\left[\log p\left(r \mid z, h\right) + \log p\left(z \mid \pi\right)\right] + const. \\
&= \sum_{n=1}^{N}\sum_{k=1}^{K} z_{nk}\left(\mathbb{E}_{q(\pi)}\left[\log \pi_k\right] + \sum_{l=1}^{L}\sum_{b\in\mathcal{B}} \mathbb{E}_{q(h)}\left[h_{klb}(r_{nlb}\log E_{nlb} + (1-r_{nlb})\log\frac{1-E_{nlb}}{\mid\mathcal{B}\mid -1})\right]\right) \\
&= \sum_{n=1}^{N}\sum_{k=1}^{K} z_{nk}\underbrace{\left(\mathbb{E}_{q(\pi)}\left[\log \pi_k\right] + \sum_{l=1}^{L}\sum_{b\in\mathcal{B}} \mathbb{E}_{q(h)}\left[h_{klb}\right]((r_{nlb}\log E_{nlb} + (1-r_{nlb})\log\frac{1-E_{nlb}}{\mid\mathcal{B}\mid -1})\right)}_{=:\log\rho_{nk}}
\end{aligned}
$$

Applying the exponential function, we derive

$$
q_x^*\left(z\right) = \prod_{n=1}^{N} q_{x_n}^*\left(z_n\right) \propto \prod_{n=1}^{N}\prod_{k=1}^{K} x_{nk}^{z_{nk}} \text{ with } x_{nk} = \frac{\rho_{nk}}{\sum_k \rho_{nk}},
$$

and

$$
\log\rho_{nk} = \mathbb{E}\left[\log\pi_k\right] + \sum_{l=1}^{L}\sum_{b\in\mathcal{B}}\mathbb{E}\left[h_{klb}\right]\left(\log {E_{nlb}}^{r_{nlb}} + \log\left(\frac{1-E_{nlb}}{\mid\mathcal{B}\mid -1}\right)^{(1-r_{nlb})}\right). \tag{2}
$$

The approximate distribution for the cluster assignments is again given by a Categorical distribution with event probabilities $(x_{nk})_{nk}$ and $\mathbb{E}_{q(z)}\left[z_{nk}\right] = x_{nk}$.

Similarly, we obtain

$$
\log q^*(\pi) = \mathbb{E}_{q(z)}\left[\log p(z \mid \pi) + \log p\left(\pi \mid \alpha\right)\right] + const.
$$

Plugging in the distributions $p(z \mid \pi)$ and $p(\pi \mid \alpha)$ and applying the exponential function, we find that the approximate distribution of the cluster weights is given by the Dirichlet distribution

$$
\begin{aligned}
q^*(\pi) = \prod_{k=1}^{K} q_{\hat{\alpha}}^*\left(\pi_k\right) \propto \prod_{k=1}^{K} &\pi_k^{\sum_n \mathbb{E}[z_{nk}]+\alpha-1} \\
&\propto Dirichlet\left(\pi \mid \sum_n \mathbb{E}\left[z_{n1}\right] + \alpha, ..., \sum_n \mathbb{E}\left[z_{nK}\right] + \alpha\right) \\
&\propto Dirichlet\left(\pi \mid \hat{\alpha}_1, ..., \hat{\alpha}_K\right),
\end{aligned}
$$

with concentration parameters $\hat{\alpha} = (\hat{\alpha}_k)_k$ where $\hat{\alpha}_k = \sum_{n=1}^{N}\mathbb{E}\left[z_{nk}\right] + \alpha$. The expected value is

$$
\mathbb{E}\left[\log\pi_k\right] = \psi(\sum_n \mathbb{E}\left[z_{nk}\right] + \alpha) - \psi(\sum_k\sum_n \mathbb{E}\left[z_{nk}\right] + \alpha)
$$

where $\psi$ is the digamma function.

For the approximate distributions of the haplotypes $q_v^*(h) = \prod_{k=1}^{K} q_{v_k}^*(h_k)$, we derive

$$
\begin{aligned}
\log q^*(h) &= \mathbb{E}\left[\log p(r \mid h, c) + \log p(h \mid \gamma)\right] + C \\
&= \sum_{k=1}^{K}\sum_{l=1}^{L}\sum_{b\in\mathcal{B}} h_{klb}\left(\log E_{nlb}\sum_{n=1}^{N} r_{nlb}\mathbb{E}\left[z_{nk}\right] + \log\frac{1-E_{nlb}}{\mid\mathcal{B}\mid -1}\sum_{n=1}^{N}(1-r_{nlb})\mathbb{E}\left[z_{nk}\right]\right) \\
&+ \sum_{k=1}^{K}\sum_{l=1}^{L}\sum_{b\in\mathcal{B}} h_{klb}\left(h_{0lb}\mathbb{E}\left[\log\gamma\right] + (1-h_{0lb})\,\mathbb{E}\left[\log\frac{1-\gamma}{\mid\mathcal{B}\mid -1}\right]\right)
\end{aligned}
$$

Defining

$$
\begin{aligned}
\log w_{klb} := \log E_{nlb}\sum_{n=1}^{N} r_{nlb}\mathbb{E}\left[z_{nk}\right] + \log\frac{1-E_{nlb}}{\mid\mathcal{B}\mid -1}\sum_{n=1}^{N}\left(1-r_{nlb}\right)\mathbb{E}\left[z_{nk}\right] \\
+ h_{0lb}\mathbb{E}\left[\log\gamma\right] + (1-h_{0lb})\,\mathbb{E}\left[\log\frac{1-\gamma}{\mid\mathcal{B}\mid -1}\right],
\end{aligned}
$$

we obtain

$$q^*(h) \propto \prod_{k=1}^{K}\prod_{l=1}^{L}\prod_{b\in\mathcal{B}} (v_{klb})^{h_{klb}}\text{ , with } v_{klb} := \frac{w_{klb}}{\sum_{b\in\mathcal{B}} w_{klb}}.$$

and the expected value is given by $\mathbb{E}[h_{klb}] = v_{klb}$.

For the approximate distribution of the mutation rate $\gamma$, we obtain

$$\begin{aligned}
&\log q^*(\gamma)\\
&= \mathbb{E}\left[\log p\left(h \mid \gamma\right) + \log p\left(\gamma_0, t_1\right)\right] + const.\\
&= \sum_k\sum_l\sum_{b\in\mathcal{B}} \mathbb{E}\left[h_{klb}\right]\left(h_{0lb}\log\gamma + (1-h_{0lb})\log\frac{1-\gamma}{\mid\mathcal{B}\mid -1}\right) + (t_0-1)\log\gamma + (t_1-1)\log 1-\gamma.
\end{aligned}$$

Applying the exponential function, we find that $q^*(\gamma) \sim Beta(\hat{t}_0, \hat{t}_1)$ with shape parameters

$$\begin{aligned}
\hat{t}_0 &= \sum_{k,l,b}\mathbb{E}\left[h_{klb}\right]h_{0lb} + t_0\\
\hat{t}_1 &= \sum_{k,l,b}\mathbb{E}\left[h_{klb}\right](1-h_{0lb}) + t_1.
\end{aligned}$$

As described above maximizing the ELBO is equivalent to minimizing the KL-divergence of the exact and the approximating distributions. Therefore the ELBO can be used as a convergence criteria for our method, and it is computed as follows:

$$\begin{aligned}
ELBO(q) =& \mathbb{E}_q\left[\log p(h,z,\gamma,\pi,r)\right] - \mathbb{E}_q\left[\log q(h,z,\gamma,\pi)\right]\\
=& \mathbb{E}_q\left[\log p(r,h)\right] + \mathbb{E}_q\left[\log p(z\mid\pi)\right] - \mathbb{E}_q\left[\log q(z)\right] + \mathbb{E}_q\left[\log p(\pi\mid\alpha)\right] - \mathbb{E}_q\left[\log q(\pi)\right]\\
&+ \mathbb{E}_q\left[\log p(h\mid\gamma)\right] - \mathbb{E}_q\left[\log q(h)\right] + \mathbb{E}_q\left[\log p(\gamma_0,t_1)\right] - \mathbb{E}_q\left[\log q(\gamma)\right]\\
=& \sum_{n=1}^{N}\sum_{k=1}^{K}\mathbb{E}_q[z_{nk}]\left(\sum_{l=1}^{L}\sum_{b\in\mathcal{B}}\mathbb{E}_q[h_{klb}]\left(r_{nlb}\log E_{nlb} + (1-r_{nlb})\log\left(\frac{1-E_{nlb}}{\mid\mathcal{B}\mid -1}\right)\right)\right)\\
&+ \sum_{n=1}^{N}\sum_{k=1}^{K}\mathbb{E}_q\left[z_{nk}\right]\mathbb{E}_q\left[\log\pi_k\right] - \sum_{k=1}^{K}\left(\log\frac{\Gamma(\hat{\alpha}_k)}{\Gamma(\sum_{k=1}^{K}\hat{\alpha}_k)} + (\hat{\alpha}_k-1)\mathbb{E}_q\left[\log\pi_k\right]\right)\\
&+ \sum_{k=1}^{K}\log\frac{1}{B(\alpha)} + (\alpha-1)\,\mathbb{E}_q\left[\log\pi_k\right] - \sum_{k=1}^{K}\log\frac{1}{B(\hat{\alpha})} + (\hat{\alpha}_k-1)\,\mathbb{E}_q\left[\log\pi_k\right]\\
&+ \sum_{k=1}^{K}\sum_{l=1}^{L}\sum_{b\in\mathcal{B}}\mathbb{E}_q[h_{klb}]\left(h_{0lb}\mathbb{E}_q\left[\log\gamma\right] + (1-h_{0lb})\,\mathbb{E}_q\left[\log\frac{1-\gamma}{\mid\mathcal{B}\mid -1}\right]\right)\\
&- \sum_{k=1}^{K}\sum_{l=1}^{L}\sum_{b\in\mathcal{B}}\mathbb{E}_q[h_{klb}]\log\mathbb{E}_q\left[h_{klb}\right]\\
&+ \log\frac{1}{B\left(t_0,t_1\right)} + (t_0-1)\,\mathbb{E}_q\left[\log\gamma\right] + (t_1-1)\,\mathbb{E}_q\left[\log 1-\gamma\right]\\
&- \log\frac{1}{B\left(\hat{t}_0,\hat{t}_1\right)} - (\hat{t}_0-1)\mathbb{E}_q\left[\log\gamma\right] - \left(\hat{t}_1-1\right)\mathbb{E}_q\left[\log 1-\gamma\right].
\end{aligned}$$

### S2 Model with uniform error rate

In the main text, we have describe in detail the model relying on the Phred-scales sequencing scores. For situations where these scores are not available, we have developed an adapted model in which the error rate $1-\theta$ is learned in an iterative fashion from the data (Figure S2). It is assumed to be identical for all sequence positions and is drawn from a Beta distribution by accounting for the matches and mismatches between the reads and their assigned haplotype. The likelihood of observing a read $r_n$ is then given by

$$p(r_{nn}, h, \theta) = \prod_{k=1}^{K} p(r_{nn} = k, h_k, \theta)^{z_{nk}} = \prod_{k=1}^{K}\left(\prod_{l=1}^{L}\prod_{b\in\mathcal{B}}\left(\theta^{r_{klb}}(\frac{1-\theta}{\mid\mathcal{B}\mid -1})^{1-r_{klb}}\right)^{h_{klb}}\right)^{z_{nk}}.$$

We develop a mean field approximation for the posterior,

$$p\left(h, z, \theta, \gamma, \pi \mid r\right) \approx q\left(h, z, \theta, \gamma, \pi\right) = q_x(z) q_v(h) q_{\hat{\alpha}}(\pi) q_{\hat{f}_0, \hat{f}_1}(\theta) q_{\hat{t}_0, \hat{t}_1}(\gamma),$$

with $\hat{f}_0$ and $\hat{f}_1$ being the variational parameters for the approximating distribution of the error parameter.

To learn the parameters of the model, we are applying the same approach as for the original model where the quality scores are used. Only the variational distributions for the cluster assignments need to be adapted:

$$\begin{aligned}
\log q^*(z) &= \mathbb{E}_{q(h,\theta,\gamma,\pi)}[\log p(r, h, \theta) + \log p(z \mid \pi)] + const. \\
&= \sum_{n=1}^{N} \sum_{k=1}^{K} z_{nk} \left( \mathbb{E}_{q(\pi)}[\log \pi_k] + \sum_{l=1}^{L} \sum_{b \in \mathcal{B}} \mathbb{E}_{q(h,\theta)}[h_{klb}(r_{nlb} \log \theta + (1 - r_{nlb}) \log \frac{1-\theta}{\mid \mathcal{B} \mid -1})] \right) \\
&= \sum_{n=1}^{N} \sum_{k=1}^{K} z_{nk} \underbrace{\left( \mathbb{E}_{q(\pi)}[\log \pi_k] + \sum_{l=1}^{L} \sum_{b \in \mathcal{B}} \mathbb{E}_{q(h)}[h_{klb}]((r_{nlb} \mathbb{E}_{q(\theta)}[\log \theta] + (1 - r_{nlb}) \mathbb{E}_{q(\theta)}[\log \frac{1-\theta}{\mid \mathcal{B} \mid -1}]) \right)}_{=: \log \rho_{nk}}
\end{aligned}$$

Applying the exponential function, we derive

$$q_x^*(z) = \prod_{n=1}^{N} q_{x_n}^*(z_n) \propto \prod_{n=1}^{N} \prod_{k=1}^{K} x_{nk}^{z_{nk}} \text{ with } x_{nk} = \frac{\rho_{nk}}{\sum_k \rho_{nk}}$$

The approximate distribution for the cluster assignments is again given by a Categorical distribution with probabilities $x = (x_n)$ and $\mathbb{E}[z_{nk}] = x_{nk}$.

For the variational distribution of the error parameter, we obtain

$$\begin{aligned}
&\log q^*(\theta) \\
&= \mathbb{E}[\log p(r, h, \theta) + \log p(\theta \mid f_0, f_1)] + const. \\
&= \sum_{n,k,l,b} r_{nlb} \left( \log \theta \mathbb{E}[z_{nk}] \mathbb{E}[h_{klb}] + \log \frac{1-\theta}{\mid \mathcal{B} \mid -1} (1 - \mathbb{E}[h_{kl}^i]) \mathbb{E}[z_{nk}] \right) \\
&+ (f_0 - 1) \log \theta + (f_1 - 1) \log 1 - \theta
\end{aligned}$$

Applying the exponential function, we find that $q_{\hat{c},\hat{d}}^*(\theta) \sim Beta\left(\hat{c}, \hat{d}\right)$ with shape parameters

$$\begin{aligned}
\hat{f}_0 &= f_0 + \sum_{n,k,l,b} r_{klb} \mathbb{E}\left[z_{nk}\right] \mathbb{E}\left[h_{klb}\right] \\
\hat{f}_1 &= f_1 + \sum_{n,k,l,b} r_{klb} \mathbb{E}\left[z_{nk}\right] \left(1 - \mathbb{E}\left[h_{klb}\right]\right).
\end{aligned}$$

Lastly, the ELBO extends to

$$
\begin{aligned}
ELBO(q) =&\mathbb{E}_q[\log p(z,\pi,h,\gamma,\theta,r)] - \mathbb{E}_q[\log q(z,\pi,h,\gamma,\theta)] \\
=&\mathbb{E}_q[\log p(r,h)] + \mathbb{E}_q[\log p(z \mid \pi)] - \mathbb{E}_q[\log q(z)] + \mathbb{E}_q[\log p(\pi \mid \alpha)] - \mathbb{E}_q[\log q(\pi)] \\
&+ \mathbb{E}_q[\log p(h \mid \gamma)] - \mathbb{E}_q[\log q(h)] + \mathbb{E}_q[\log p(\gamma \mid t_0,t_1)] - \mathbb{E}_q[\log q(\gamma)] \\
&+ \mathbb{E}_q[\log p(\theta \mid f_0,f_1)] - \mathbb{E}_q[\log q(\theta)] \\
=&\sum_{n=1}^{N}\sum_{k=1}^{K}\mathbb{E}_q[z_{nk}]\left(\sum_{l=1}^{L}\sum_{b\in\mathcal{B}}\mathbb{E}_q[h_{klb}]\left(r_{nlb}\mathbb{E}[\log\theta] + (1-r_{nlb})\mathbb{E}\left[\log\frac{1-\theta}{\mid\mathcal{B}\mid -1}\right]\right)\right) \\
&+\sum_{n=1}^{N}\sum_{k=1}^{K}\mathbb{E}_q\left[z_{nk}\right]\mathbb{E}_q\left[\log\pi_k\right] - \sum_{k=1}^{K}\left(\log\frac{\Gamma(\hat{\alpha}_k)}{\Gamma(\sum_{k=1}^{K}\hat{\alpha}_k)} + (\hat{\alpha}_k-1)\mathbb{E}_q[\log\pi_k]\right) \\
&+\sum_{k=1}^{K}\log\frac{1}{B(\alpha)} + (\alpha_k-1)\mathbb{E}_q[\log\pi_k] - \sum_{k=1}^{K}\log\frac{1}{B(\hat{\alpha})} + (\hat{\alpha}_k-1)\mathbb{E}_q[\log\pi_k] \\
&+\sum_{k=1}^{K}\sum_{l=1}^{L}\sum_{b\in\mathcal{B}}\mathbb{E}_q[h_{klb}]\left(h_{0lb}\mathbb{E}_q[\log\gamma] + (1-h_{0lb})\mathbb{E}_q\left[\log\frac{1-\gamma}{\mid\mathcal{B}\mid -1}\right]\right) \\
&-\sum_{k=1}^{K}\sum_{l=1}^{L}\sum_{b\in\mathcal{B}}\mathbb{E}_q[h_{klb}]\log\mathbb{E}_q[h_{klb}] \\
&+\log\frac{1}{B(t_1,t_2)} + (t_1-1)\mathbb{E}_q\left[\log\gamma\right] + (t_1-1)\mathbb{E}_q\left[\log 1-\gamma\right] \\
&-\log\frac{1}{B\left(\hat{t}_0,\hat{t}_1\right)} + (\hat{t}_0-1)\mathbb{E}_q[\log\gamma] + (\hat{t}_1-1)\mathbb{E}_q\left[\log 1-\gamma\right] \\
&+\log\frac{1}{B(f_0,f_1)} + (f_0-1)\,\mathbb{E}_q\left[\log\theta\right] + (f_1-1)\mathbb{E}_q\left[\log 1-\theta\right] \\
&-\log\frac{1}{B(\hat{f}_0,\hat{f}_1)} + (\hat{f}_0-1)\mathbb{E}_q\left[\log\theta\right] + (\hat{f}_1-1)\mathbb{E}_q\left[\log 1-\theta\right].
\end{aligned}
$$

### S3 VILOCA-*envp*: Exclusion of non-variable positions in the clustering

As discussed in the main text Section ”Methods”, VILOCA’s inference algorithm is scaling linearly with the number of reads (coverage) $N$ and size of the local region $L$. In order to reduce the runtime, we developed a method where users can decide to exclude positions of low or no variation. Users can adjust the *envp*-threshold. By setting this threshold, any positions where the fraction of reads that do not match the reference base is below the threshold will be excluded from the clustering step. At those excluded positions, the reference base is reported. Hence mutations of frequency below the *envp*-threshold cannot be detected.

Let $M \subset \{1,\dots,L\}$ be the set of positions where all reads match the reference $h_0$, i.e.:

$$\forall l \in M, \forall i \in \mathcal{A} : h^i_{0,l} = r^i_{n,l}, \text{ for all } n \in \{1,\dots,N\}.$$

Then, in those ”matching” positions, we assume that all reconstructed haplotypes coincide with the master sequence and reads:

$$\forall l \in M, \forall k \in \{1,\dots,K\} : h^i_{k,l} = h^i_{0,l}, \text{ for all } i \in \mathcal{A}.$$

Let $M^c = M \setminus \{1,\dots,L\}$ denote the complement set of $M$, the set of variable positions, meaning the positions where not all reads match the reference.

We show that the set of matching positions contributes only with a constant term to the ELBO. We first consider the standard model using quality scores. The likelihood for read $r_n$ is given by

$$
\begin{aligned}
p(r_n \mid z_n, h) &= \prod_k\left[\prod_l\prod_b\left(E_{n,l}^{r_{nlb}}\left(\frac{1-E_{nl}}{\mid\mathcal{B}\mid -1}\right)^{1-r_{nlb}}\right)^{h_{klb}}\right]^{z_{nk}} \\
&= \prod_k\left[\prod_{l\in M^c}\prod_b(\dots)^{h_{klb}}\right]^{z_{nk}}\prod_k\left[\prod_{l\in M}\prod_b\left(E_{n,l}^{r_{nlb}}\left(\frac{1-E_{n,l}}{\mid\mathcal{B}\mid -1}\right)^{1-r_{nlb}}\right)^{h_{klb}}\right]^{z_{nk}}.
\end{aligned}
$$

For any $l \in M$, if $r_{nlb} = 1$, then $h_{klb} = 1$, hence

$$(E_{n,l})^{r_{nlb}} \left( \frac{1 - E_{n,l}}{\mid \mathcal{B} \mid -1} \right)^{1 - r_{nlb}} = E_{nlb},$$

else, if $r_{nlb} = 0$, then $h_{klb} = 0$, and hence

$$(E_{nlb})^{r_{nlb}} \left( \frac{1 - E_{nlb}}{\mid \mathcal{B} \mid -1} \right)^{1 - r_{nlb}} = 1.$$

It follows that

$$\begin{aligned} p(r_n \mid z_n, h) &= \prod_k \left[ \prod_{l \in M^c} \prod_b \left( E_{n,l}^{r_{nlb}} \left( \frac{1 - E_{nl}}{\mid \mathcal{B} \mid -1} \right)^{1 - r_{nlb}} \right)^{h_{klb}} \right]^{z_{nk}} \prod_k \left[ \prod_{l \in M} E_{nl} \right]^{z_{nk}} \\ &= \prod_k \left[ \prod_{l \in M^c} \prod_b \left( E_{n,l}^{r_{nlb}} \left( \frac{1 - E_{nl}}{\mid \mathcal{B} \mid -1} \right)^{1 - r_{nlb}} \right)^{h_{klb}} \right]^{z_{nk}} \underbrace{\prod_{l \in M} E_{nl}}_{= const.} . \end{aligned}$$

Hence, we find that considering only the set of mismatching positions changes the read likelihood only by a constant factor. Correspondingly, the matching positions also contribute only with a constant term to the ELBO. To show this, we are only concerned with the terms marked with $(*)$ as the others are not depending on the genome positions:

$$\begin{aligned} \text{ELBO}(q) = & \underbrace{\mathbb{E}\left[\log p\left(r \mid z, h\right)\right]}_{(*)} + \mathbb{E}\left[\log p\left(z \mid \pi\right)\right] - \mathbb{E}\left[\log q\left(z\right)\right] + \mathbb{E}\left[\log p\left(\pi \mid \alpha\right)\right] - \mathbb{E}\left[\log q\left(\pi\right)\right] \\ & + \underbrace{\mathbb{E}\left[\log p\left(h \mid \gamma\right)\right] - \mathbb{E}\left[\log q\left(h\right)\right]}_{(*)} + \underbrace{\mathbb{E}\left[\log p\left(\gamma, b\right)\right] - \mathbb{E}\left[\log q\left(\gamma\right)\right]}_{(*)} . \end{aligned}$$

For the first term $\mathbb{E}[\log p(r \mid z, h)]$, define $C_{nlb} := \log E_{nlb} r_{nlb}$. Then,

$$\begin{aligned} \mathbb{E}\left[\log p\left(r, h\right)\right] &= \sum_n \sum_k \mathbb{E}\left[z_{nk}\right] \left( \sum_{l \in M^c} \cdots \right) + \sum_n \sum_k \mathbb{E}\left[z_{nk}\right] \sum_{l \in M} \sum_{i \in \mathcal{A}} C_{nlb} \\ &= \sum_n \sum_k \mathbb{E}\left[z_{nk}\right] \left( \sum_{l \in M^c} \cdots \right) + \sum_n \sum_{l \in M} \sum_{i \in \mathcal{A}} C_{nlb} \underbrace{\left( \sum_k \mathbb{E}\left[z_{nk}\right] \right)}_{=1 \text{ by construction}} \\ &= \sum_n \sum_k \mathbb{E}\left[z_{nk}\right] \left( \sum_{l \in M^c} \cdots \right) + \underbrace{\sum_n \sum_{l \in M} \sum_{i \in \mathcal{A}} C_{nlb}}_{= const.} . \end{aligned}$$

For the second term, we can show that the matching positions only contribute with a constant to the ELBO:

$$\begin{aligned} & \mathbb{E}\left[\log p\left(h \mid \gamma\right)\right] - \mathbb{E}\left[\log q\left(h\right)\right] \\ = & \sum_k \left( \sum_{l \in M^c} \sum_{b \in \mathcal{B}} \cdots + \sum_{l \in M} \sum_{b \in \mathcal{B}} \underbrace{\underbrace{\mathbb{E}\left[h_{klb}\right]}_{=1} \left( \underbrace{h_{0lb}}_{=1} \mathbb{E}\left[\log \gamma\right] + \underbrace{(1 - h_{0lb})}_{=0} \mathbb{E}\left[\log \frac{1 - \gamma}{\mid \mathcal{A} \mid -1}\right] \right)}_{=1(1\mathbb{E}[\log \gamma])} \right) \\ & - \sum_k \left( \sum_{l \in M^c} \cdots + \underbrace{\sum_{l \in M} \sum_{i \in \mathcal{A}} \mathbb{E}\left[h_{klb}\right] \log \mathbb{E}\left[h_{klb}\right]}_{=0, \text{ if } i \text{ s.t. } h_{0lb}=1, \text{ then } \mathbb{E}[h_{klb}]=1, \text{ else } \mathbb{E}[h_{klb}]=0} \right) \\ = & \sum_k \left( \sum_{l \in M^c} \sum_{b \in \mathcal{B}} \cdots + \sum_{l \in M} \mathbb{E}\left[\log \gamma\right] \right) - \sum_k \left( \sum_{l \in M^c} \cdots + \sum_{l \in M} 0 \right) \end{aligned}$$

Lastly, we consider the term of the mutation rate:

$$\mathbb{E}\left[\log p(\gamma \mid a, b)\right] - \mathbb{E}\left[\log q(\gamma)\right] = \log \frac{B(\hat{a}, \hat{b})}{B(a, b)} + (a - 1 - \hat{a} + 1)\mathbb{E}\left[\log \gamma\right] + \left(b - 1 - \hat{b} + 1\right) \mathbb{E}\left[\log 1 - \gamma\right],$$

where

$$\hat{a} = \sum_k \sum_{l \in M^c} \sum_{b \in \mathcal{B}} \cdots + a + \sum_k \sum_{l \in M} \underbrace{\sum_{b \in \mathcal{B}} \mathbb{E}\left[h_{klb}\right] h_{0lb}}_{=1} = \sum_k \sum_{l \in M^c} \sum_{b \in \mathcal{B}} \cdots + a + K \mid M \mid$$

$$\hat{b} = \sum_k \sum_{l \in M^c} \sum_{b \in \mathcal{B}} \cdots + b + \sum_k \sum_{l \in M} \underbrace{\sum_{b \in \mathcal{B}} \mathbb{E}\left[h_{klb}\right] (1 - h_{0lb})}_{=1} = \sum_k \sum_{l \in M^c} \sum_{b \in \mathcal{B}} \cdots + b.$$

In summary, the matching positions are only contributing with constant factors to the ELBO.

For the model in which an error rate is learned from the data, the derivations are very similar. With same reasoning as before, the likelihood is given by:

$$p(r_n \mid z_n, h, \theta) = \prod_k \left[ \prod_{l \in M^c} \prod_{b \in \mathcal{B}} (\dots)^{h_{klb}} \right]^{z_{nk}} \prod_{l \in M} \theta = \theta^{|M|} \prod_k \left[ \prod_{l \in M^c} \prod_{b \in \mathcal{B}} (\dots)^{h_{klb}} \right]^{z_{nk}}$$

To check the effect of the matching positions on the ELBO, we need to only be concerned with the terms marked with $(*)$ as the others are not depending on the genome positions:

$$\begin{aligned} \mathrm{ELBO}(q) = & \underbrace{\mathbb{E}\left[\log p\left(r \mid z, h, \theta\right)\right]}_{(*)} + \mathbb{E}\left[\log p\left(z \mid \pi\right)\right] - \mathbb{E}\left[\log q\left(z\right)\right] + \mathbb{E}\left[\log p\left(\pi \mid \alpha\right)\right] - \mathbb{E}\left[\log q\left(\pi\right)\right] \\ & + \underbrace{\mathbb{E}\left[\log p\left(h \mid \gamma\right)\right] - \mathbb{E}\left[\log q\left(h\right)\right]}_{(*)} + \underbrace{\mathbb{E}\left[\log p\left(\gamma, b\right)\right] - \mathbb{E}\left[\log q\left(\gamma\right)\right]}_{(*)} + \underbrace{\mathbb{E}\left[\log p\left(\theta, d\right)\right] - \mathbb{E}\left[\log q\left(\theta\right)\right]}_{(*)}. \end{aligned}$$

For the first term, we get

$$\begin{aligned} \mathbb{E}\left[\log p\left(r \mid z, h, \theta\right)\right] &= \sum_{n,k} \mathbb{E}\left[z_{nk}\right] \left( \sum_{l \in M^c} \cdots \right) \\ &\quad + \sum_{n,k} \mathbb{E}\left[z_{nk}\right] \left( \sum_{l \in M} \sum_{b \in \mathcal{B}} \mathbb{E}\left[h_{klb}\right] \left( r_{nlb} \mathbb{E}\left[\log \theta\right] + (1 - r_{nlb})\, \mathbb{E}\left[\log \frac{1 - \theta}{\mid \mathcal{B} \mid -1}\right] \right) \right) \\ &= \sum_{n,k} \mathbb{E}\left[z_{nk}\right] \left( \sum_{l \in M^c} \cdots \right) + \sum_n \underbrace{\sum_k \mathbb{E}\left[z_{nk}\right]}_{=1} \mid M \mid \mathbb{E}\left[\log \theta\right] \\ &= \sum_{n,k} \mathbb{E}\left[z_{nk}\right] \left( \sum_{l \in M^c} \cdots \right) + N \mid M \mid \mathbb{E}\left[\log \theta\right]. \end{aligned}$$

The terms $\mathbb{E}\left[\log p(h \mid \gamma)\right] - \mathbb{E}\left[\log q(h)\right]$ and $\mathbb{E}\left[\log p(\gamma \mid t_0, t_1)\right] - \mathbb{E}[\log q(\gamma)]$ solve as for the quality score model. Lastly,

$$\begin{aligned} & \mathbb{E}\left[\log p\left(\theta \mid f_0, f_1\right)\right] - \mathbb{E}\left[\log q(\theta)\right] \\ & = \log \frac{B\left(\hat{f}_0, \hat{f}_1\right)}{B\left(f_0, f_1\right)} + \left(f_{-}1 - \hat{f}_0 + 1\right) \mathbb{E}\left[\log \theta\right] + \left(f_0 - 1 - \hat{f}_1 + 1\right) \mathbb{E}\left[\log 1 - \theta\right], \end{aligned}$$

where

$$\hat{c} = \sum_{n,k} \sum_{l \in M^c} \sum_{b \in \mathcal{B}} \cdots + c + \sum_{n,k} \mathbb{E}\left[z_{nk}\right] \sum_{l \in M} \underbrace{\sum_{b \in \mathcal{B}} r_{nlb} \mathbb{E}\left[h_{klb}\right]}_{=1} = \sum_{n,k} \mathbb{E}\left[z_{nk}\right] \sum_{l \in M^c} \sum_{b \in \mathcal{B}} \cdots + c + N \mid M \mid$$

$$\hat{d} = \sum_{n,k} \mathbb{E}\left[z_{nk}\right] \sum_{l \in M^c} \sum_{b \in \mathcal{B}} \cdots + d + \sum_{n,k} \mathbb{E}\left[z_{nk}\right] \sum_{l \in M} \underbrace{\sum_{b \in \mathcal{B}} r_{nlb} (1 - \mathbb{E}\left[h_{klb}\right])}_{=0} = \sum_{n,k} \mathbb{E}\left[z_{nk}\right] \sum_{l \in M^c} \sum_{b \in \mathcal{B}} \cdots + d.$$

To conclude, also for the model with error rate parameter the matching positions only contribute to the ELBO by a constant shift.

### S4 Benchmarking study

In this section, we provide additional details on our benchmarking study. The complete benchmarking study can be reproduced using the Snakemake workflow provided on GitHub (`https://github.com/cbg-ethz/V-pipe/tree/feature-benchmark/resources/auxiliary_workflows/benchmark/resources/local_haplotype_setup`). It was conducted using the benchmarking module of V-pipe 3.0 [2].

#### S4.1 Reproducing the benchmarking study

To reproduce the benchmarking study and create the figures presented in the manuscript, use the following instructions:

1. Clone the repository of V-pipe 3.0 into your working directory:
   ```
   git clone https://github.com/cbg-ethz/V-pipe.git
   ```
2. Go into the directory of the benchmarking study for the local haplotype reconstruction:
   ```
   cd V-pipe/resources/auxiliary_workflows/benchmark/resources/local_haplo_setup
   ```
3. The parameters to reproduce the synthetic datasets are in the parameter files `config_xxx/params.csv` with the configuration file `config_xxx/config.yaml` where simulation mode, replicate number and methods to be executed are defined.
4. The methods to execute must be define in a Python script in this directory:
   ```
   V-pipe/resources/auxiliary_workflows/benchmark/resources/method_definitions.
   ```
5. Now the workflow is ready, go back to the directory
   ```
   cd V-pipe/resources/auxiliary_workflows/benchmark/resources/local_haplo_setup
   ```
6. To install the needed Conda environments execute:
   ```
   snakemake --conda-create-envs-only --use-conda -c1
   ```
7. To submit the workflow to a lsf-cluster execute `./run_workflow.sh`, otherwise execute the workflow with `snakemake --use-conda -c1`.
8. The workflow will provide the results in the directory `results`.
9. When the workflow has terminated and all result files were generated, figures from the manuscript can be generated by executing the notebooks in `./workflow/notebooks/`.

#### S4.2 Failure of CliqueSNV in processing the simulated long-read samples.

CliqueSNV encountered difficulties in achieving convergence for the simulated PacBio samples. Initially, we attempted running CliqueSNV with the parameter *-tf=0.001*, but it failed to converge within the 14-day and 23-hour runtime limit that was set due to available computational resources. The documentation of the tool (`https://github.com/vtsyvina/CliqueSNV`) mentions that using a low value for *-tf* can significantly increase the runtime and memory requirements. As a result, we also tried executing CliqueSNV with *-tf=0.01* and *-tf=0.1*. However, even with this adjustment, the tool still did not achieve convergence and only provided a consensus sequence. The runtime of CliqueSNV is quadratic with respect to the number of SNVs which is likely to cause the significantly runtime increase. It should be noted however that CliqueSNV generally tries to solve the more difficult problem of global haplotype reconstruction, where reads covering different regions are being connected.

### S5 Applications

In this section, we provide additional details on the data processing of the samples presented in the main text section Applications.

#### S5.1 HIV-1 Illumina samples

We analysed 9 longitudinal samples from an HIV-positive patient (accession numbers in Supplementary Table S2) . We used the tool V-pipe 3.0 [2] for quality control and read alignment of the samples. For the read alignment we used the alignment tool bwa-mem with default settings. Subsequently, we applied VILOCA with uniform tiling strategy with local regions size of 300 base pairs as this was the reported approximate mean read length across the samples. Additionally, we omitted regions that where only covered by less than 20 reads (*–win_coverage 20*). To ensure high confident mutation calls we set the posterior score cutoff to 0.9. As a comparison, we also applied the mutation calling tool LoFreq version 2.1.5 [3] with default parameters. To process each sample VILOCA took on average 57 minutes with a maximal memory usage of 903 MB. The full analysis and figure can be reproduced using the Snakemake workflow and the notebook available on GitHub (`https://github.com/cbg-ethz/viloca_applications/tree/main/hiv_clinical`).

#### S5.2 SARS-CoV-2 Illumina wastewater samples from Zurich

We analysed 56 wastewater samples that were collected in the scope of the Swiss SARS-CoV-2 Wastewater Surveillance efforts ([4]; Accession numbers in Supplementary Table SS3). The samples were processed using the data processing tool V-pipe 3.0 [2], using iVar trim [5] for primer trimming, and the tool bwa mem [6] to align sequencing reads to the reference (NC_045512.2). VILOCA was applied using the amplicon-tiling strategy, where the local regions were defined based on the ARTIC v4 amplicon protocol. For each amplicon, we defined two local regions of 250bp length that are overlapping in the middle of the amplicon. The full analysis and figure can be reproduced using the Snakemake workflow available on GitHub (`https://github.com/cbg-ethz/SARS-CoV-2-wastewater-sample-processing-VILOCA/tree/main/resources/setup_emergence_new_variant`).

#### S5.3 MinIon wastewater samples from Pakistan

We analysed a sewage sample from Pakistan (accession number: ERR4033236) that was taken in the scope of the official polio surveillance program by the WHO and Pakistan Government [7]. The sample was processed using the data processing tool V-pipe 3.0 [2]. We used the alignment tool minimap2 [8] to align reads to the Human poliovirus 1 reference genome (AY560657.1). As the vast majority of the aligned reads spanned most parts of the VP1 region, VILOCA was applied to a single genomic region spanning the entire VP1 gene. To process the sample VILOCA took 19 minutes with a maximal memory usage of 1635 MB. The complete scripts that were used to analyse the sample are publicly available on GitHub (`https://github.com/cbg-ethz/viloca_applications/tree/main/polio_sewage`).

Table S1: Configurations used for the generation of distance based haplotype populations. $n_1$: number of haplotypes in group one; $n_2$: number of haplotypes in group two; $d_{12}$: average pairwise distance between group one and two; $d_1$: average pairwise sequence distance within group one; $d_2$: average pairwise sequence distance within group two. We generated 5 replicates for each haplotype population configuration.

| Population | Sequencing technology | Genome size | Coverage | Number of haplotypes | Haplotype frequencies | Pairwise distances | $n_1$ | $n_2$ | $d_{12}$ | $d_1$ | $d_2$ |
|---|---|---|---|---|---|---|---|---|---|---|---|
| 1 | Illumina | 249 | 1000 | 5 | 33%, 25%, 18%, 14%, 10% | 8.0-12.0 % | 2 | 3 | 30 | 20 | 20 |
| 2 | Illumina | 249 | 100, 200, 500, 800, 1000, 5000, 10000 | 10 | 26%, 20%, 15%, 11%, 8%, 6%, 5%, 4% , 3%, 2% | 4.0-12.0 % | 5 | 5 | 30 | 10 | 15 |
| 3 | Illumina | 249 | 1000 | 15 | 25%, 19%, 14%, 11%, 8%, 6%, 5%, 3%, 2.5%, 2%, 1.4%, 1.1%, 0.8%, 0.6%, 0.5% | 4.0-12.0 % | 5 | 10 | 30 | 10 | 10 |
| 4 | Illumina | 249 | 1000 | 20 | 25%, 19%, 14%, 11%, 8%, 6%, 4%, 3%, 2.5%, 2%, 1.4%, 1%, 0.8%, 0.6%, 0.4%, 0.3%, 0.25%, 0.2%, 0.14% 0.1% | 4.0-12.0 % | 5 | 15 | 30 | 10 | 15 |
| 5 | ONT, PacBio | 5000 | 1000 | 5 | 33%, 25%, 18%, 14%, 10% | 8-12.0 % | 2 | 3 | 600 | 400 | 400 |
| 6 | ONT, PacBio | 5000 | 100, 200, 500, 800, 1000, 5000, 10000 | 5 | 33%, 25%, 18%, 14%, 10% | 12.0 % | 2 | 3 | 600 | 600 | 600 |
| 7 | ONT, PacBio | 5000 | 1000 | 10 | 26%, 20%, 15%, 11%, 8%, 6%, 5%, 4% , 3%, 2% | 4.0-12.0 % | 5 | 5 | 600 | 200 | 300 |
| 8 | ONT, PacBio | 5000 | 1000 | 15 | 25%, 19%, 14%, 11%, 8%, 6%, 5%, 3%, 2.5%, 2% , 1.4%, 1.1%, 0.8%, 0.6%, 0.5% | 4.0-12.0 % | 5 | 10 | 600 | 200 | 240 |

Initialize.<br>
**while** $ELBO(q)$ has not converged **do**<br>
    **for** $n \in \{1, ..., N\}$ and $k \in \{1, ..., K\}$ **do**<br>
        update $\mathbb{E}[z_{nk}]$<br>
    **end for**<br>
    **for** $k \in \{1, ..., K\}$, $l \in \{1, ..., L\}$ and $i \in \{1, ..., B\}$ **do**<br>
        update $\mathbb{E}[h_{klb}]$<br>
    **end for**<br>
    **for** $k \in \{1, ..., K\}$ **do**<br>
        update $\mathbb{E}[\log \pi_k]$<br>
    **end for**<br>
    update $\mathbb{E}[\log \gamma], \mathbb{E}[\log 1 - \gamma]$<br>
Compute $ELBO(q)$<br>
**end while**

Figure S1: Coordinate ascent variational inference algorithm. Through iterative updates, the parameters defining the variational distributions in the mean field approximation are learned. In each iteration, the expected value of each factorized variational distribution is updated while keeping the other parameters fixed. The update equations for each expected value are detailed in the Supplementary Material section Model using the sequencing quality scores. Convergence is evaluated by monitoring the absolute changes in the Evidence Lower Bound (ELBO). The algorithm terminates if the absolute change falls below the convergence threshold (default: 1e-03).

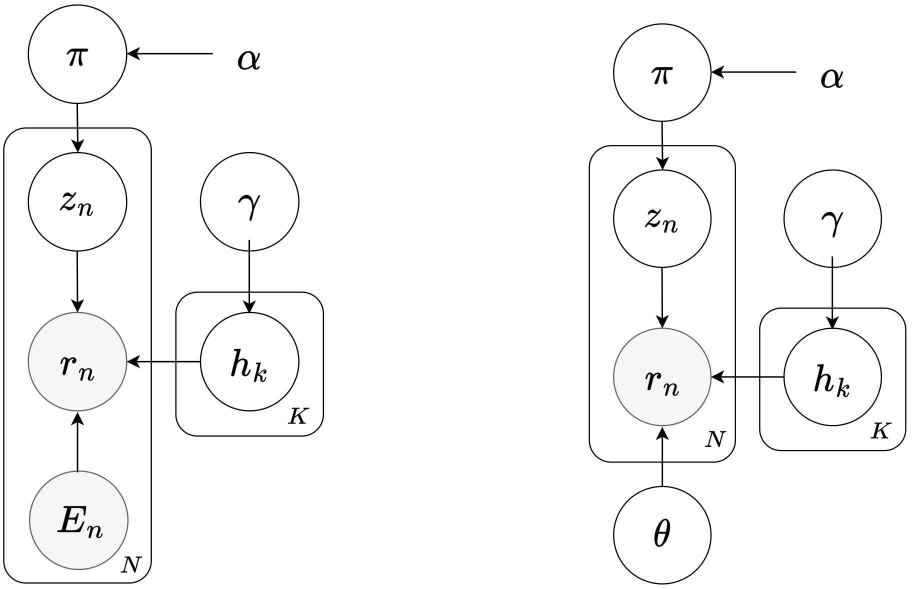


(a) Quality score model.

(b) Model with latent error parameter.

Figure S2: Graphical representation of the Mixture Models: each node represents a random variable, shading denotes the observed variables, plates denote replications of the model, the number of replications is given in the bottom right corner.

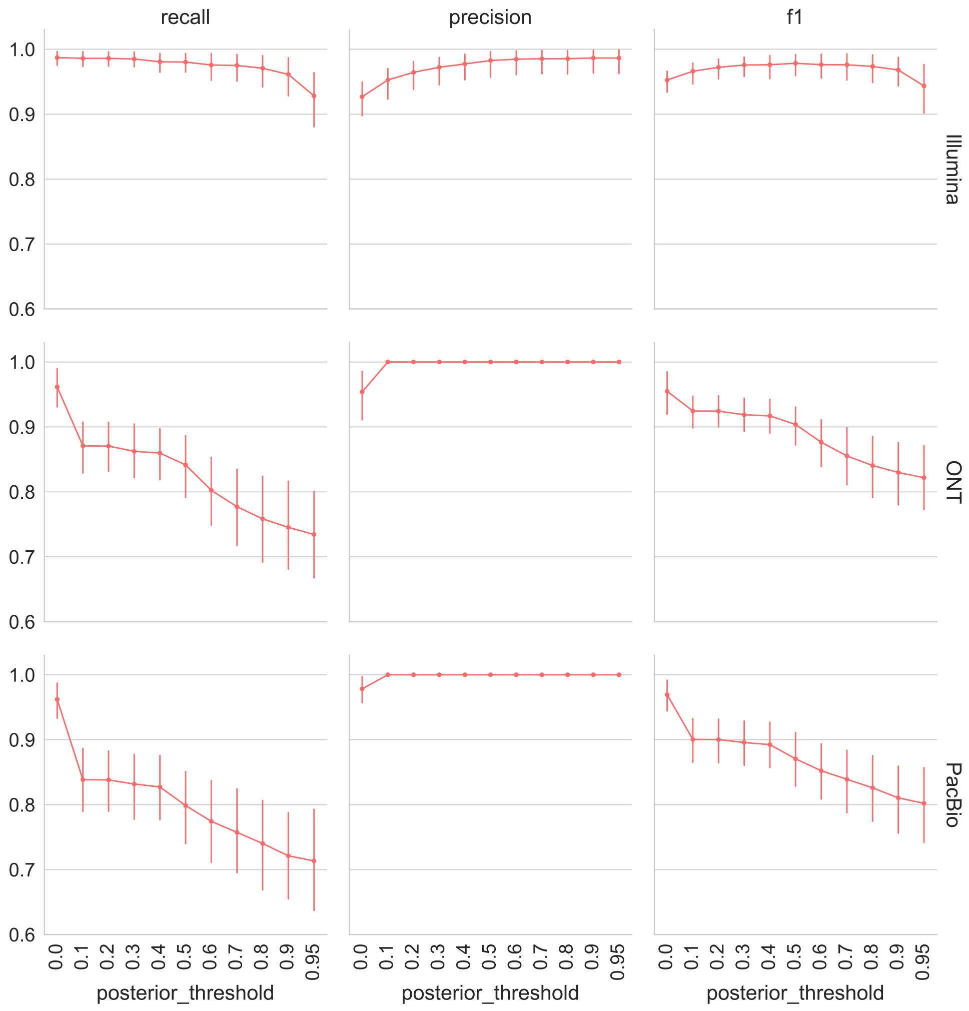


Figure S3: VILOCA mutation calling performance in terms of precision, recall and f1-score for varying posterior thresholds on all simulated Illumina, ONT, and PacBio samples as defined in Table S1.

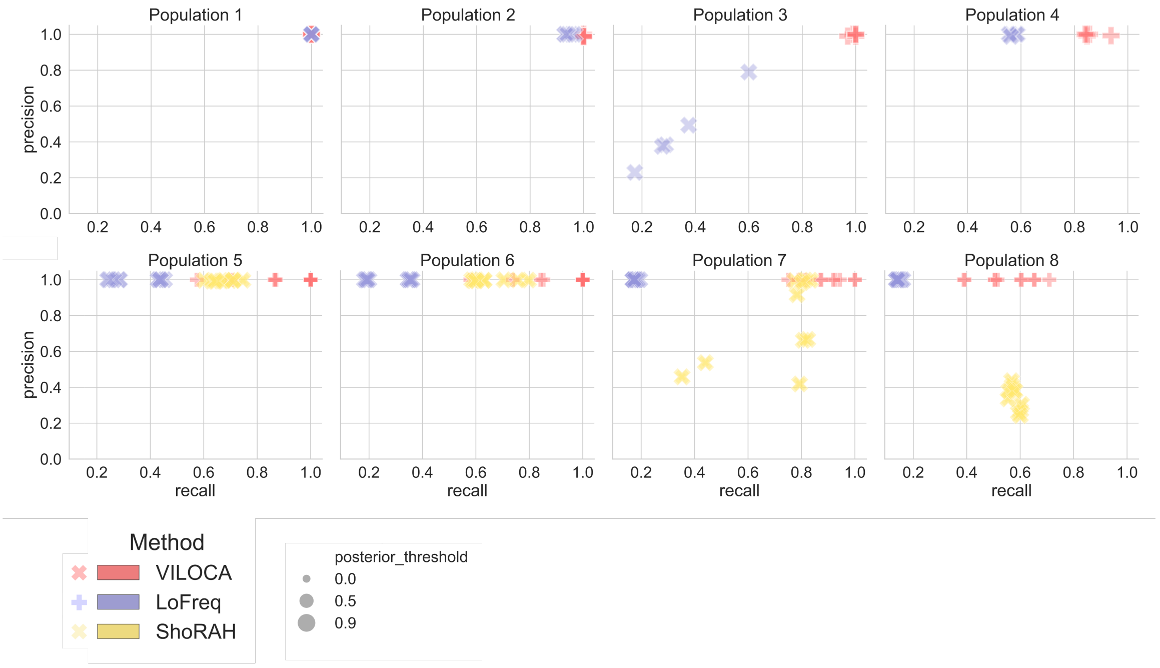


Figure S4: Performance evaluation in terms of mutation calling for simulated long- and short-read samples. Precision-recall plots for each simulated population with short-read samples displayed in the upper row and long-read samples in the lower row both with coverage 1000. VILOCA and ShoRAH provide confidence scores for each mutation call, represented by posterior distributions. Marker sizes indicate the posterior threshold utilized for mutation calls, while LoFreq results are represented by single-sized markers.

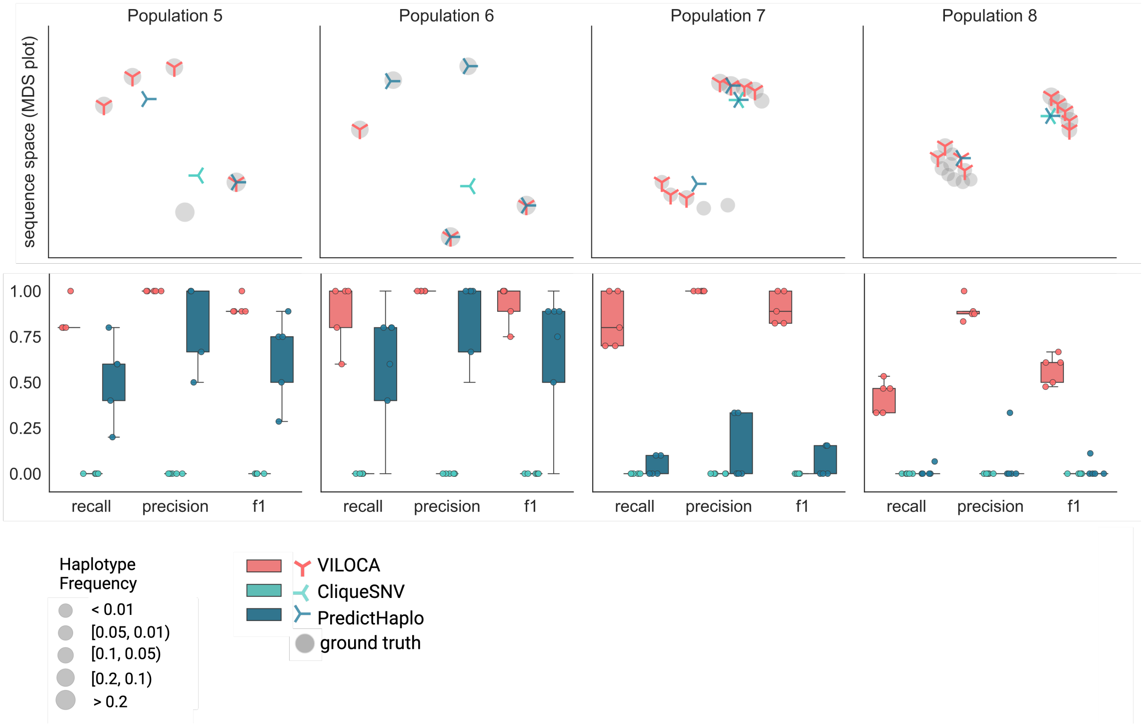


Figure S5: Performance on simulated ONT samples. Upper row: MDS plots for the sequence space visualization of one example simulation replicate per haplotype population. Each marker represents one sequence. Symbol size corresponds to the frequency of the respective haplotype in the sample. Lower row: Precision, recall and F1-score for respective population across the five replicates. For the computation of true and false positives, we employ a distance threshold of 0.01 between prediction and ground truth sequence.

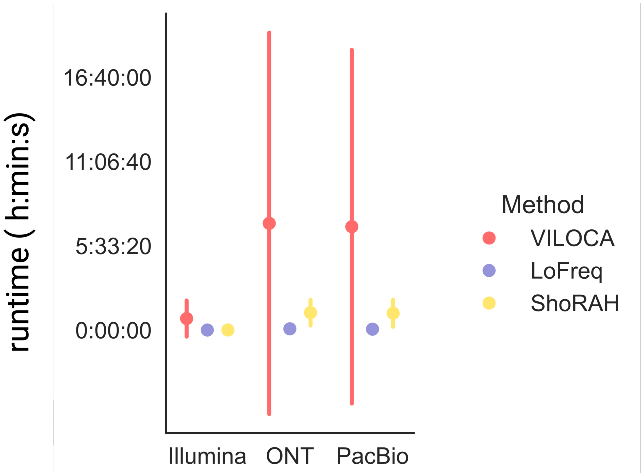


Figure S6: Runtime comparison of the thee methods for mutation calling across all simulated datasets of Illumina, ONT, and PacBio reads. Error bars are depicting the standard deviation.

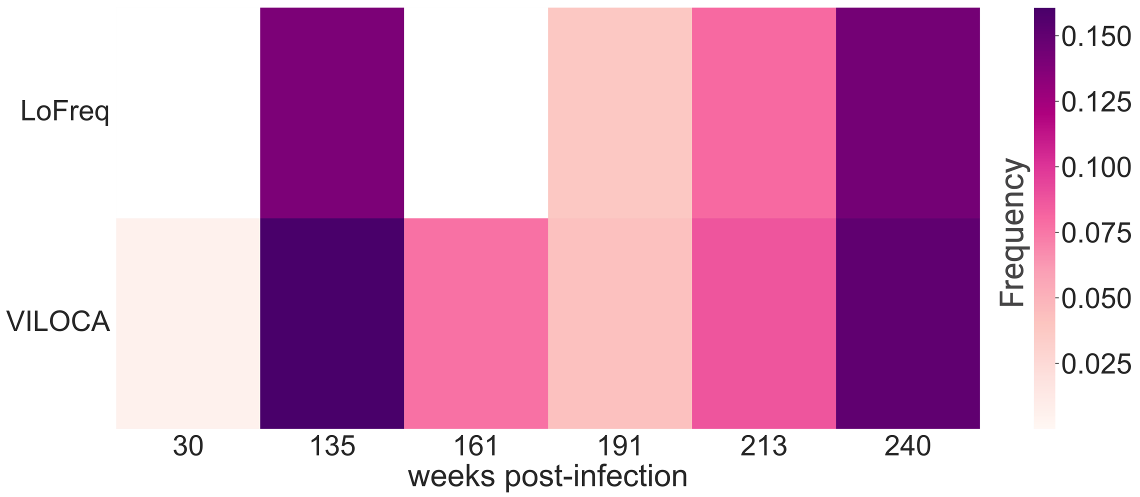


Figure S7: Application of VILOCA to longitudinal Illumina samples of an HIV-1 positive patient pre-treatment. Heatmap of prevalence predictions by VILOCA and LoFreq of mutation C982T in the longitudinal samples.

Table S2: Details to the Illumina samples of the HIV-positive patient with patient-id CAP257. Mean read coverage were assessed with V-pipe [2].

| Accession number | Week post-infection | Mean coverage |
|---|---|---|
| SRR9588783 | 30 | 597 |
| SRR9588794 | 135 | 274 |
| SRR9588793 | 161 | 411 |
| SRR9588792 | 191 | 1887 |
| SRR9588791 | 213 | 899 |
| SRR9588790 | 240 | 1464 |

Table S3: Table of the ENA accession numbers analysed SARS-CoV-2 Illumina wastewater samples from Zurich.

| Sample name | ENA sample accession number |
|---|---|
| A2_10_2022_04_14 | ERS18825839 |
| A2_10_2022_04_21 | ERS18825838 |
| A2_10_2022_04_28 | ERS18826529 |
| A2_10_2022_05_05 | ERS18826528 |
| A2_10_2022_05_12 | ERS18826527 |
| A2_10_2022_06_02 | ERS18826553 |
| A2_10_2022_06_09 | ERS18825828 |
| B2_10_2022_04_15 | ERS18826526 |
| B2_10_2022_04_22 | ERS18826525 |
| B2_10_2022_04_29 | ERS18825808 |
| B2_10_2022_05_06 | ERS18825807 |
| B2_10_2022_05_13 | ERS18825806 |
| B2_10_2022_06_03 | ERS18826559 |
| B2_10_2022_06_10 | ERS18825840 |
| B2_10_2022_06_17 | ERS18826431 |
| B2_10_2022_06_24 | ERS18707709 |
| C2_10_2022_04_16 | ERS18825805 |
| C2_10_2022_04_23 | ERS18825804 |
| C2_10_2022_04_30 | ERS18825803 |
| C2_10_2022_05_07 | ERS18825802 |
| C2_10_2022_05_14 | ERS18825801 |
| C2_10_2022_05_30 | ERS18826537 |
| C2_10_2022_06_04 | ERS18826565 |
| C2_10_2022_06_11 | ERS18825846 |
| C2_10_2022_06_25 | ERS18707715 |
| C6_10_2022_05_16 | ERS18826513 |
| D2_10_2022_04_17 | ERS18826512 |
| D2_10_2022_04_24 | ERS18826511 |
| D2_10_2022_05_01 | ERS18826510 |
| D2_10_2022_05_08 | ERS18826509 |
| D2_10_2022_05_15 | ERS18826508 |
| D2_10_2022_06_05 | ERS18826572 |
| D2_10_2022_06_12 | ERS18825851 |
| D2_10_2022_06_26 | ERS18707720 |
| E2_10_2022_04_18 | ERS18825800 |
| E2_10_2022_04_25 | ERS18825799 |
| E2_10_2022_05_02 | ERS18825798 |
| E2_10_2022_05_09 | ERS18826507 |
| E2_10_2022_06_06 | ERS18825786 |
| E2_10_2022_06_06 | ERS18825786 |
| E2_10_2022_06_13 | ERS18825856 |
| E2_10_2022_06_20 | ERS18826471 |
| F2_10_2022_04_12 | ERS18826506 |
| F2_10_2022_04_19 | ERS18826505 |
| F2_10_2022_04_26 | ERS18826504 |
| F2_10_2022_05_03 | ERS18826503 |
| F2_10_2022_05_10 | ERS18826502 |
| F2_10_2022_06_07 | ERS18825791 |
| F2_10_2022_06_14 | ERS18825902 |
| F2_10_2022_06_21 | ERS18826476 |
| H1_10_2022_04_13 | ERS18826501 |
| H1_10_2022_04_20 | ERS18826500 |
| H1_10_2022_04_27 | ERS18826499 |
| H1_10_2022_05_04 | ERS18826498 |
| H1_10_2022_05_11 | ERS18826497 |
| H1_10_2022_05_18 | ERS18826496 |
| H1_10_2022_06_01 | ERS18825819 |
| H1_10_2022_06_08 | ERS18826430 |
| H1_10_2022_06_15 | ERS18826485 |
